# Cooperativity of catalytic and lectin-like domain of *T. congolense* trans-sialidase modulates its catalytic activity

**DOI:** 10.1101/2021.05.28.446113

**Authors:** Mario Waespy, Thaddeus T. Gbem, Nilima Dinesh Kumar, Shanmugam Solaiyappan Mani, Jana Rosenau, Frank Dietz, Sørge Kelm

**Author notes:** Corresponding Author: Dr. Mario Waespy, Centre for Biomolecular Interactions Bremen, Faculty for Biology and Chemistry, University Bremen, Bremen, 28359, Germany.

## Abstract

Trans-sialidases (TS) represent a multi-gene family of unusual enzymes, which catalyse the transfer of terminal sialic acids from sialoglycoconjugates to terminal galactose or *N*-acetylgalactosamine residues of oligosaccharides without the requirement of CMP-Neu5Ac, the activated Sia used by typical sialyltransferases. Most work on trypanosomal TS has been done on enzymatic activities of TS from *T. cruzi* (causing Chagas disease in Latin America), subspecies of *T. brucei*, (causing human sleeping sickness in Africa) and *T. congolense* (causing African Animal Trypanosomosis in livestock). Previously, we demonstrated that *T. congolense* TS (TconTS) lectin domain (LD) binds to several carbohydrates, such as 1,4-β-mannotriose.

To investigate the influence of TconTS-LD on enzyme activities, we firstly performed *in silico* analysis on structure models of TconTS enzymes. Findings strongly supports the potential of domain swaps between TconTS without structural disruptions of the enzymes overall topologies. Recombinant domain swapped TconTS1a/TS3 showed clear sialidase and sialic acid (Sia) transfer activities, when using fetuin and lactose as Sia donor and acceptor substrates, respectively. While Sia transfer activity remained unchanged from the level of TconTS1a, hydrolysis was drastically reduced. Presence of 1,4-β-mannotriose during TS reactions modulates enzyme activities favouring trans-sialylation over hydrolysis.

In summary, this study provides strong evidence that TconTS-LDs play pivotal roles in modulating enzyme activity and biological functions of these and possibly other TS, revising our fundamental understanding of TS modulation and diversity.

## Introduction

Trypanosomes are protozoan parasites causing trypanosomiasis in Southern America (caused by *Trypanosoma cruzi*), also known as Chagas’ disease (see review by Clayton 2010 (Clayton 2010)), Human African Trypanosomiasis (HAT, caused by *Trypanosoma brucei ssp*.) and Animal African Trypanosomiasis (AAT, also called Nagana caused by *Trypansoma congolense*) in livestock in Sub-Saharan Africa. AAT brings death to millions of cattle annually (World Health Organization 2013; Kamuanga 2003). To evade insect and mammalian host immune systems, parasites developmental stage specifically express unusual enzymes termed trans-sialidases (TS). TS catalyse the transfer of terminal sialic acids (Sia) from host glycoconjugates to terminal galactose residues on target glycoproteins (Schenkman et al. 1991; Engstler et al. 1995; Buschiazzo et al. 2002). Several studies have shown that trypanosomal TS play important roles in the pathology of the disease in mammalian host (Schenkman et al. 1991; Montagna et al. 2002; Nok & Balogun 2003; Coustou et al. 2012). Structurally, all currently known trypanosomal TS contain two domains, a N-terminal catalytic domain (CD) and a C-terminal lectin-like domain (LD), which are connected via a 23 to 25 amino acid long α-helix (Buschiazzo et al. 2002).

Whereas published studies have focused on the enzymatic activities and catalytic mechanism of TS-CDs (Campetella et al. 1994; Cremona et al. 1995; París et al. 2001; Haselhorst 2004; Koliwer-Brandl et al. 2011; Oliveira et al. 2014), no experimental biological function of the LDs has been described so far.

Smith and Eichinger reported the expression and characterisation of *Trypanosoma cruzi* TS (TcruTS)/ *Trypanosoma rangeli* sialidase (TranSA) hybrid proteins, exhibiting different Sia transfer and sialidase activities (Smith & Eichinger 1997). They found that the C-terminal Fn3 domain (fibronectin type III), named according to its structural relation to fibronectin type III, is not only required for expression of enzymatically active TcruTS and TranSA (Pereira et al. 1991; Schenkman et al. 1994; Smith et al. 1996), but also influences the overall Sia transfer and sialidase activities (Smith & Eichinger 1997). Amino acid sequence alignments of a well characterised sialidase from *M. viridifaciens* (Gaskell et al. 1995) with TcruTS revealed that R572 and E578, which are known to be essential for galactose binding of *M. viridifaciens* sialidase (Gaskell et al. 1995), are well conserved in TcruTS and TranSA (Smith & Eichinger 1997). Point mutation of one of these residues resulted in reduced sialidase activities for both enzymes and enhanced Sia transfer activity in TcruTS (Smith & Eichinger 1997). As a consequence of these findings Smith and Eichinger predicted both amino acid residues (R572 and E578 in TcruTS) to be involved in galactose binding of the acceptor and/or donor substrates, which would necessarily require an overall protein folding that brings the catalytic domain and the region containing the Arg and Glu of Fn3 domain (at least R and E) close together (Smith & Eichinger 1997). However, the resolved crystal structures of TcruTS (Buschiazzo et al. 2002) and TranSA (Buschiazzo et al. 2000; Amaya et al. 2003) demonstrated that these two amino acid residues are located far away from the actual Sia binding pocket of the catalytic domain, and therefore, they are unlikely to be directly involved in substrate binding as proposed by Smith and Eichinger (Smith & Eichinger 1997). Nevertheless, even if these residues do not interact with the galactose moiety of the acceptor/donor substrate, R611 and E617 of TranSA (correspond to R572 and E578 in TcruTS) were found to form an intramolecular salt bridge at the C-terminus of the LD (Amaya et al. 2003) apparently indirectly influencing enzymatic activities, since its disruption was found to change Sia transfer and sialidase activities, respectively (Smith & Eichinger 1997). Interestingly, structural and amino acid sequence alignments of TcruTS (EMBL: AAA66352.1), *Tryoanosoma brucei* (Tbru) TS (EMBL: AAG32055.1), TranSA (EMBL: AAC95493.1), *Trypanosoma vivax* (Tviv) TS (CCD20961.1) and *Trypanosoma congolense* (Tcon) TS revealed that this salt bridge is conserved among these TS (data not shown), indicating a possibly essential role in trypanosomal trans-sialidase activities. Surprisingly, to the best of our knowledge no further investigations regarding functional data of the TS-LD were reported until now.

First evidence for a more pivotal role of the LDs has come from our phylogenetic analysis done separately on CD and LD of TconTS (Gbem et al. 2013). Previously, we demonstrated the binding of TconTS-LD to oligomannose and oligogalactose oligosaccharides (Waespy et al. 2015). Interestingly, mannose and oligomannose oligosaccharides are not Sia acceptor substrates for the catalytic transfer (Engstler et al. 1993; Engstler et al. 1995; Tiralongo et al. 2003). However, oligomannose oligosaccharides have been found in *N*- and *O*-linked glycans on glycoproteins or as part of their glycosylphosphatidylinositol (GPI)-anchor on the parasite’s surface (Savage et al. 1984; Zamze et al. 1990; Zamze et al. 1991; Bayne et al. 1993; Beecroft et al. 1993; Bütikofer et al. 2002; Thomson et al. 2002). Therefore, these glycans potentially function as ligand structures for TconTS-LD. Furthermore, also TS were found to be glycosylated, predominantly with *N*-linked glycans of the high-mannose type (Engstler & Schauer 1993; Pontes de Carvalho et al. 1993), leading to the suggestion of intermolecular interactions possibly mediated by TS-LD. Evidence for this has come from experiments demonstrating the mannose-dependent oligomerisation of recombinant high-mannosylated TconTS (Waespy et al. 2015).

Tiralongo *et al*. prepared the anti-TconTS monoclonal antibody (mAb) 7/23, which recognises two TconTS forms isolated from procyclic *T. congolense* culture supernatant (Tiralongo et al. 2003). Later, T. Gbem *et al*. showed the specific binding of anti-TconTS mAb 7/23 to recombinant TconTS1 and TconTS2, purified from cell culture supernatants of transfected CHO-Lec1 cells. Recombinant TconTS3 and TconTS4 were not recognised by the antibody (Gbem et al. 2013). However, the epitope of this antibody has not been identified so far.

Here we report a strategy to effectively swap CDs and LDs from different TconTS in order to elucidate the functional relationship between these domains and to investigate a potential impact of LDs in modulating enzymatic activity. Homology model of TconTS revealed that amino acid residues localised at the contact sites between TconTS CD and LD are well conserved in the TS family. Furthermore, *in silico* data of domain-swapped TconTS revealed a similar overall topology with an extensive hydrogen bond network at the interface between CD and LD. Based on these observations we assume that swapping of LDs from different TconTS would not interfere with overall structural arrangements (Koliwer-Brandl et al. 2011). Enzymatic activity data of bacterial- and CHO-Lec1-expressed TconTS1a and domain-swapped TconTS1a/TS3 demonstrated that the LD directly affects the overall catalytic activity of the enzyme. The existence and strength of a simultaneous and cooperative binding of CD and LD to sialic acid and oligomannose structures of the same substrate respectively, is decisive for enzymatic activity. Reducing or blocking the binding of TconTS1a-LD either by swapping the LD of TconTS1a with that of TconTS3 exhibiting no affinity to oligomannose structures (Waespy et al. 2015) or by using 1,4β-mannotriose as a competitive inhibitor during reactions changed the trans-sialidase over sialidase activity (TS/sialidase) ratio significantly. In summary, our results clearly demonstrate the involvement of TconTS-LD in modulating catalytic activities of TconTS.

## Results

### The contact sites between TconTS CD and LD

Using the crystal structure of *T. cruzi* TS (TcruTS) (Buschiazzo et al. 2002) as template, homology models of TconTS1-4 were calculated as described under Methods. Similar to other TS, CD and LD are localised in close proximity, connected by a 23 to 25 amino acid long α-helix (Figure 1).

**Figure 1:**
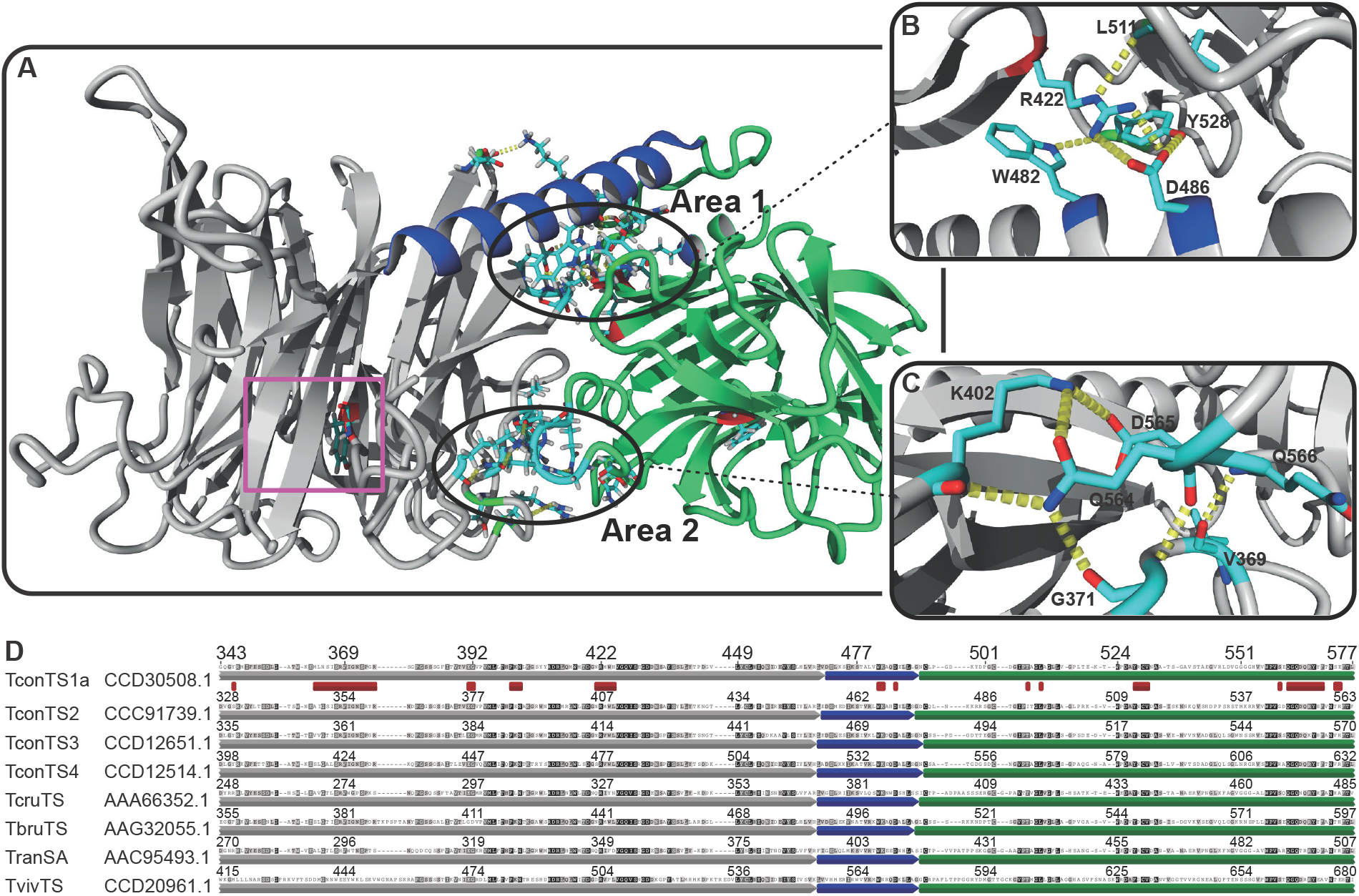
Hydrogen bond network at the interface between CD and LD of TconTS. **A:** homology model of TconTS1a using the crystal structure of TcruTS (PDB code: 3B69) as template structure as described under Methods. Area 1 and 2, comprising a network of hydrogen bonds formed by amino acid residues at the interface between catalytic domain (CD in grey), interdomain α-helix (blue) and lectin domain (LD in green) are marked. The active centre containing the catalytic tyrosine residue Tyr438 at the CD is labelled with a purple square **B-C:** Zoom of Area 1 and Area 2 showing the conserved hydrogen bond network (yellow dotted lines), respectively. **D:** Amino acid sequence alignment (see methods for details) of TconTS1a through TconTS4, *T. cruzi* TS (TcruTS EMBL: AAA66352.1), *T. brucei* TS (TbruTS EMBL: AAG32055.1), *T. rangeli* sialidase (TranSA EMBL: AAC95493.1) and *T. vivax* TS (TvivTS EMBL: CCD20961.1). Only a section of the complete alignment is shown including contact sites between CD and LD. CD is labelled in grey, α-helix in blue and LD in green respectively. Sequence segments being part of the interface between CD, α-helix and LD are marked with red squares. Increasing background darkness for each residue of the sequence indicates increasing number of identical amino acid residues at the corresponding position over the alignment (see methods for details).

From the homology models, it was observed that the arrangement of CD and LD is stabilised by close contact sites between both. The interface between CD and LD from TcruTS (Buschiazzo et al. 2002) and *T. rangeli* sialidase (TranSA) (Amaya et al. 2003), both South American species, were determined by X-ray crystallography. Both were found to be about 36 to 48 % larger compared to those from sialidases with a lectin-like domain, such as *Vibrio cholerae* sialidase (VCS) (Crennell et al. 1994) and *Marcobdella decora* (leech) intramolecular trans-sialidase (IT-sialidase) (Luo et al. 1999). Along this line, molecular interfaces of several African TS including TconTS1, TconTS2, TconTS3, TconTS4, *T. brucei* TS (TbruTS) and *T. vivax* TS (TvivTS) were determined from homology structure models (see Methods for details). Results revealed 33 to 46 % larger areas compared to that of VCS and leech IT-sialidase for all African TS, consistent with findings for the South American species as shown in Table 1. It can be seen that the overall surface area of the contact sites between CD and LD of African TS is around 2833.55 to 3306.47 Å^2^ and quite similar in size among the trypanosomal TS family. A detailed *in silico* analysis addressing potential interactions between amino acids at the contact sites of TconTS, revealed a network of hydrogen bonds. This network is expected to stabilise a comparative rigid overall conformation of the enzyme. Notably, calculated structure models revealed two defined locations, in which the majority of inter-domain hydrogen bonds were concentrated (Figure 1).

**Table 1:**
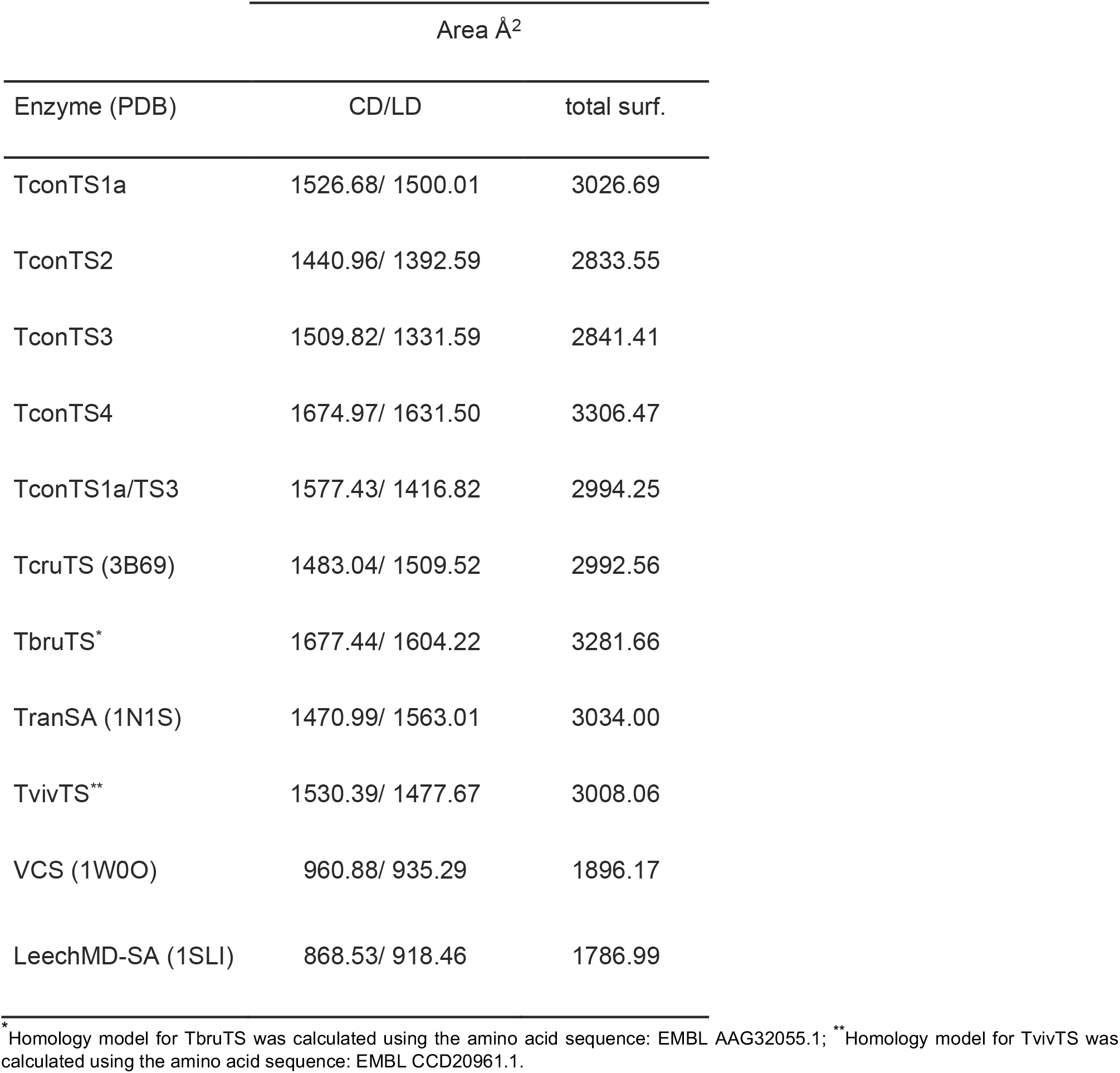
Calculated molecular surface of the contact site between CD and LD of trypanosomal TSs and common sialidases.

One site (Area 1) is closer to the α-helix connecting both domains (Figure 1A and B), whereas the second (Area 2) is located opposite of area 1 (Figure 1A and C). Amino acids of both domains forming inter-domain hydrogen bonds at the CD/LD interface of TconTS1 through TconTS4 are summarised in Table 2.

**Table 2:**
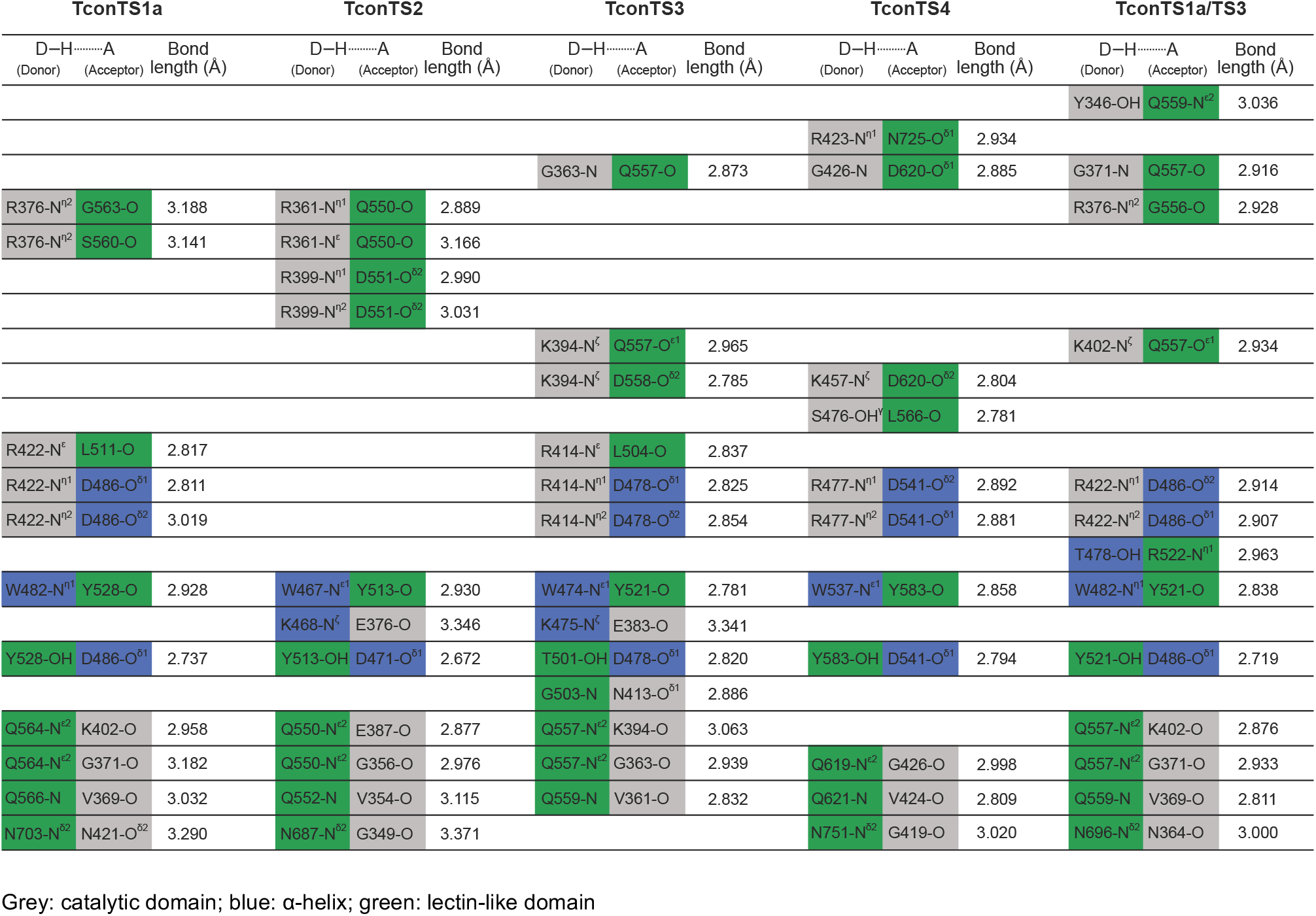
Hydrogen bonds formed by amino acid residues at the contact site between CD and LD of TconTS

Energy minimised homology models revealed 13 hydrogen bonds formed at the interface between CD, LD and the α-helix for TconTS3, 12 for TconTS1, and 11 for both, TconTS2 and TconTS4. Moreover, the number of these hydrogen bonds in each TS are equally distributed between both areas 1 and 2 (Figure 1B and C). Not surprisingly, amino acid sequence alignments of TS revealed that amino acid residues essential for hydrogen bond formation in area 1 and 2 are well conserved among the TS family (Table 3). For example, TconTS and TbruTS are highly conserved in these contact areas among each other with no more than three amino acid variations from the consensus sequence (Table 3). In contrast, TvivTS shows 10 amino acid changes relative to the consensus sequence. This is not surprising since it has been shown that TvivTS is more distant related to TconTS and TbruTS (Gbem et al. 2013).

**Table 3:** Conserved amino acid residues involved in hydrogen bond formation between CD and LD at the contact site of TS

|  | sn | Consensus | TconTS1a | TconTS2 | TconTS3 | TconTS4 | TbruTS* | TvivTS** | TcruTS*** | TranSA**** |
| --- | --- | --- | --- | --- | --- | --- | --- | --- | --- | --- |
| **CD** | 1 | Gly | N364 | G349 | G356 | G419 | G376 | G439 | G269 | G291 |
|  | 2 | Arg | R368 | R353 | R360 | R423 | R380 | E443 | R273 | H295 |
|  | 3 | Val | V369 | V354 | V361 | V424 | V381 | E443 | V274 | V296 |
|  | 4 | Gly | G371 | G356 | G363 | G426 | G383 | W446 | G276 | T298 |
|  | 5 | Arg | R376 | R361 | R368 | R431 | R388 | V451 | S281 | S303 |
|  | 6 | Glu | E391 | E376 | E383 | E446 | G410 | E473 | E296 | E318 |
|  | 7 | Lys | K402 | E387 | K394 | K457 | K421 | K484 | L307 | L329 |
|  | 8 | Asn | N421 | N406 | N413 | S476 | N440 | D503 | Q326 | Q348 |
|  | 9 | Arg | R422 | R407 | R414 | R477 | R441 | R504 | R327 | R349 |
| **αHel** | 10 | Trp | W482 | W467 | W474 | W537 | W501 | W569 | W386 | W408 |
|  | 11 | Lys | K483 | K468 | K475 | K538 | K502 | N570 | K387 | K409 |
|  | 12 | Asp | D486 | D471 | D478 | D541 | D505 | D573 | D390 | D412 |
| **LD** | 13 | Thr | T508 | I493 | T501 | T563 | T528 | T604 | T417 | T439 |
|  | 14 | Leu | L511 | L496 | L504 | L566 | L531 | L607 | L420 | L442 |
|  | 15 | Tyr | Y528 | Y513 | Y521 | Y583 | Y548 | F629 | Y437 | Y459 |
|  | 16 | Ser | S560 | G546 | S553 | R615 | S580 | S663 | S468 | A490 |
|  | 17 | Gly | G563 | G549 | G556 | G618 | G583 | G666 | G471 | G493 |
|  | 18 | Gln | Q564 | Q550 | Q557 | Q619 | Q584 | Q667 | Q472 | Q494 |
|  | 19 | Asp | D565 | D551 | D558 | D620 | D585 | D668 | N473 | T495 |
|  | 20 | Gln | Q566 | Q552 | Q559 | Q621 | Q586 | R669 | Q474 | R496 |
|  | 21 | Asn | N674 | N658 | N667 | N725 | H694 | A778 | D581 | D603 |

\* TbruTS sequence: EMBL AAG32055.1; \*\* TtivTS sequence: EMBL CCD20961.1; \*\*\* TcruTS sequence: EMBL AAA66352.1; \*\*\*\* TranSA sequence: EMBL AAC95493.1. It should be noted that different TbruTS have been identified, which group together with TconTS1, 3 and 4, respectively (Gbem et al. 2013). sn: serial number.

With 9 deviations from the consensus, most amino acid changes were found in TranSA, 5 in CD and 4 in LD (Table 3). Of these, 4 are identical in TcruTS, which exhibits 5 changes in total, 3 in CD and 2 in LD.

The three amino acid residues Trp serial number (sn) 10, Lys sn 11 and Asp sn 12 of the α-helix are essential for the formation of hydrogen bonds to Glu sn 6 and Arg sn 9 of CD and Tyr sn 15 and Trp sn 13 (Table 3) of LD in Area 1 (Figure 1B). Interestingly, these amino acids are conserved through all TS as listed in Table 3 except TvivTS, in which Lys sn 11 is replayced by an Asn residue, indicating a fundamental role of that region for enzyme structure preservation. In addition, it can be seen that the indole ring of Trp sn 10 provides additional van der Waals interactions with the aliphatic side chain of Arg sn 9 similar to such an interaction observed in Siglec-1 (Sialoadhesin) (Zaccai et al. 2003). The Gln sn 18 of LD, which is conserved in the trypanosomal TS family, is located on a loop at the more exposed Area 2 (Figure 1C). It reaches relatively deep into the CD, where it forms hydrogen bonds to Gly sn 4, Lys sn 7 and Arg sn 5. In summary, the relatively large interface between the CD and LD of trypanosomal TS compared to related sialidases together with the extended hydrogen bond network formed by well-conserved amino acids within the interdomain interface appears to stabilise a distinct orientation of both domains relative to each other.

### TconTS domain swap

To investigate the influence of TconTS-LD on enzyme activities, we established a strategy to swap CDs and LDs from different TconTS. A structure model of domain swap TS TconTS1a/TS3 was calculated *in silico* and found to predict a similar overall topology for such a recombinant TS as in the models for TconTS1a and TconTS3 (Figure 2).

**Figure 2:**
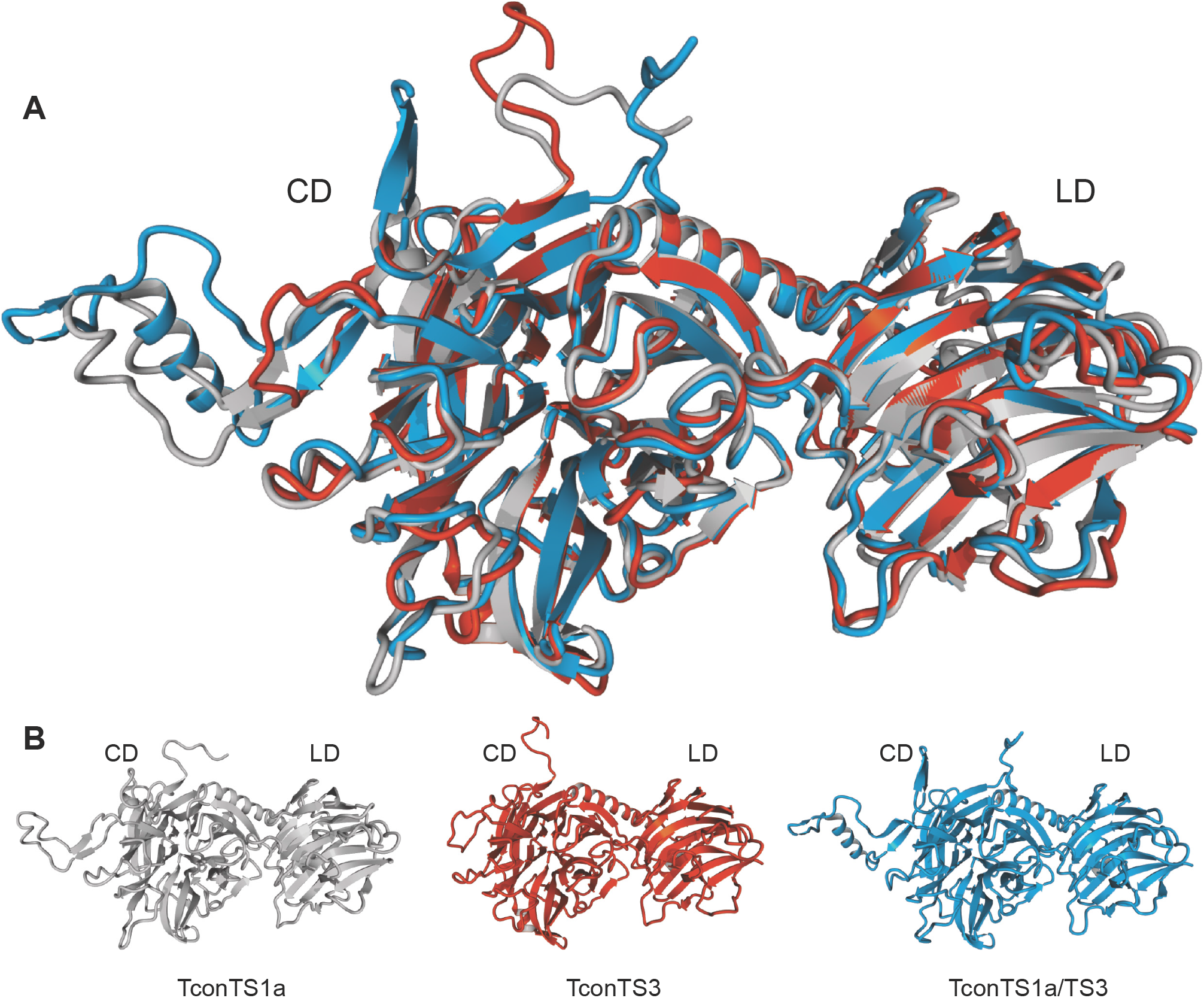
Homology model of domain swapped TconTS1a/TS3. **A:** Structural alignment of TconTS1a, TconTS3 and domain swapped TconTS1a/TS3 homology model. **B:** Homology model of TconTS1a (grey), TconTS3 (red) and TconTS1a/TS3 (blue). Catalytic domain (CD) and lectin-like domain (LD) are indicated. Homology models and structure alignment were generated using the YASARA *Structure* module as described under methods.

Crystal structure analysis of TcruTS (Buschiazzo et al. 2002) revealed a crystallographically unresolved flexible loop right after the α-helix between the two domains (Figure 3). This structural hairpin loop is stabilised by a well-conserved disulphide bridge formed between Cys493 and Cys503 in TconTS1a (Figure 3A). Besides these two cysteine residues, no conserved amino acids are found in this region of the four TconTS, which even differ in length (Figure 3A). Based on these observations we hypothesised that this loop region can tolerate a wide range of mutations and thus would be suitable for the fusion of CDs and LDs from different TconTS. Therefore, we introduced an Eco105I (SnaBI) endonuclease restriction site (coding for the dipeptide Tyr-Val) between the codes for D497 and K498 of TconTS1a and the corresponding positions in the other TconTS (Figure 3B). This strategy facilitated the possibility for convenient fusion of any LD to any CD in order to swap the entire LDs, without disrupting any potentially important structure elements such as β-sheets, salt-bridges or α-helices in the enzyme.

**Figure 3:**
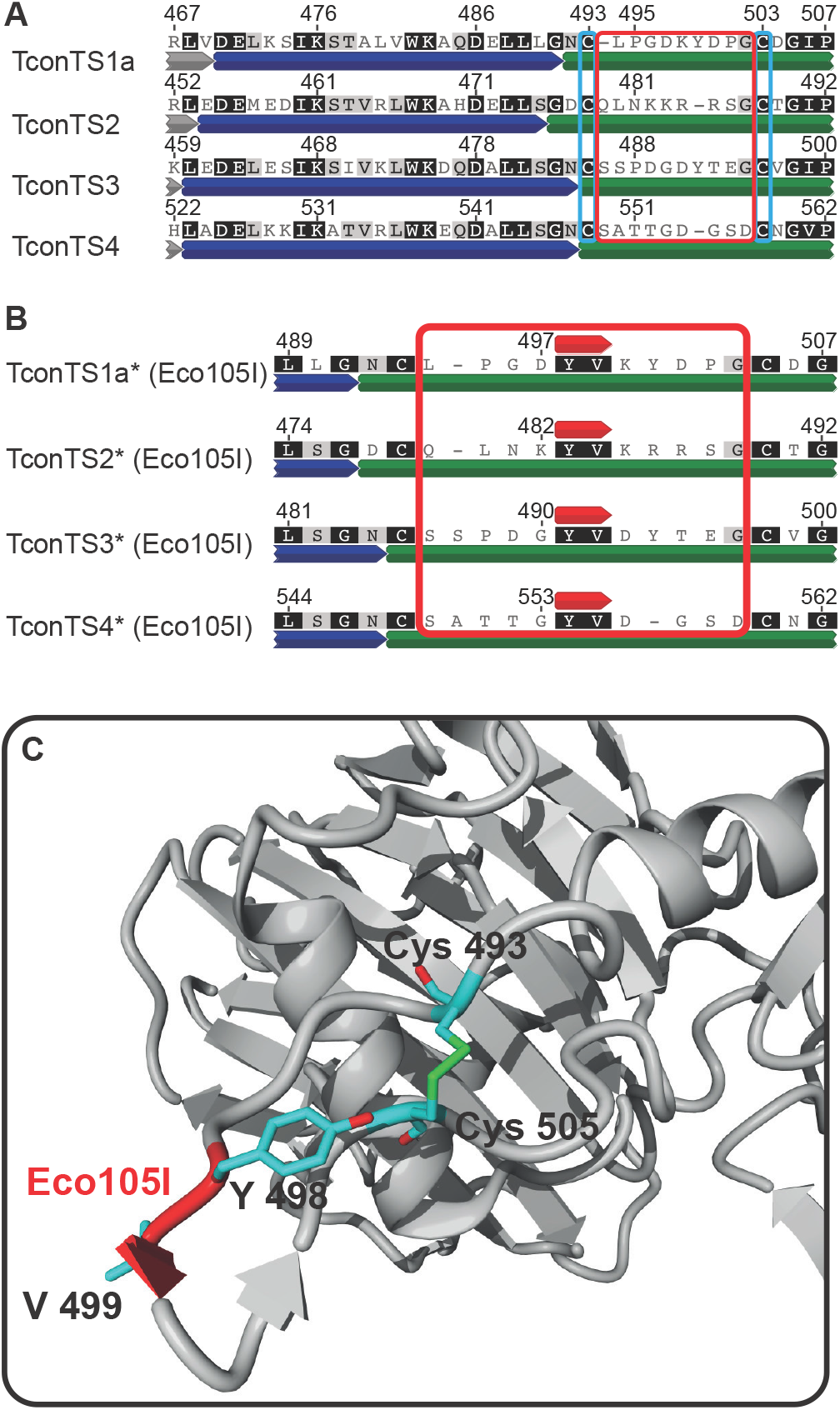
Insertion of Eco105I endonuclease restriction site into TconTS. **A:** A section of the complete amino acid sequence alignment of TconTS1a through TconTS4 with assigned structural elements (Grey, blue and green for CD, α-helix and LD, respectively). Cysteine residues Cys493 and Cys503 in TconTS1a forming a well conserved disulphide bridge in TS are marked with light blue frames, whereas the resulting hairpin loop flanked by these cysteine residues is marked with a red frame. **B:** Amino acid sequence alignment of the hairpin loop region with the Eco105I restriction site inserted (red annotation above each TconTS sequence, resulting in Tyr-Val insertion). Alignments were calculated as described under Methods. Numbers on top of each sequence indicate the corresponding residue numbers in the individual TconTS before the insertion. TconTS comprising the inserted Eco105I restriction site are labelled (*). Increasing darkness of background for each residue indicates increasing number of identical amino acid residues at the corresponding position. **C:** Homology model of TconTS1a* showing a region of the lectin domain (grey), including the well conserved disulphide bridge Cys493-Cys505 and the hairpin loop with the Eco105I insertion (red label). TconTS1a* homology model was calculated as described under Methods.

### Expression and purification of domain swapped TconTS

Recombinant TconTS were expressed in prokaryotic (pTconTS) and eukaryotic (eTconTS) cells. All four recombinant pTconTS* (TconTS containing *Eco*105I restriction site, Table 4) were expressed by *E. coli* Rosetta (DE3) pLacI, as described under Methods. A schematic illustration of expressed recombinant pTconTS*, is shown in Figure 4B. Sufficient amounts of pTconTS* were obtained and characterised by SDS-PAGE and Western blot analysis revealing molecular masses of about 135 kDa (Figure 4C). However, only relatively low amounts (100 – 300 μg of enzyme per litre of bacterial culture) of soluble pTconTS* were obtained, whereas the majority of the recombinant proteins were insoluble. Several expression optimisations, including variation of the isopropyl-β-D-1-thiogalactopyranoside (IPTG) concentration, as well as time of induction and temperature adjustments slightly increased yields to about 200 – 500 μg of purified soluble protein per litre of bacterial culture. In the light of previous studies (Koliwer-Brandl et al. 2011; Gbem et al. 2013; Waespy et al. 2015), it has become apparent that LDs are implicated in the enzyme activity of TconTS. Therefore, domain swap of LDs from a highly and a less active TconTS (e.g. TconTS1 and TconTS3) represent a logic target model system.

**Figure 4:**
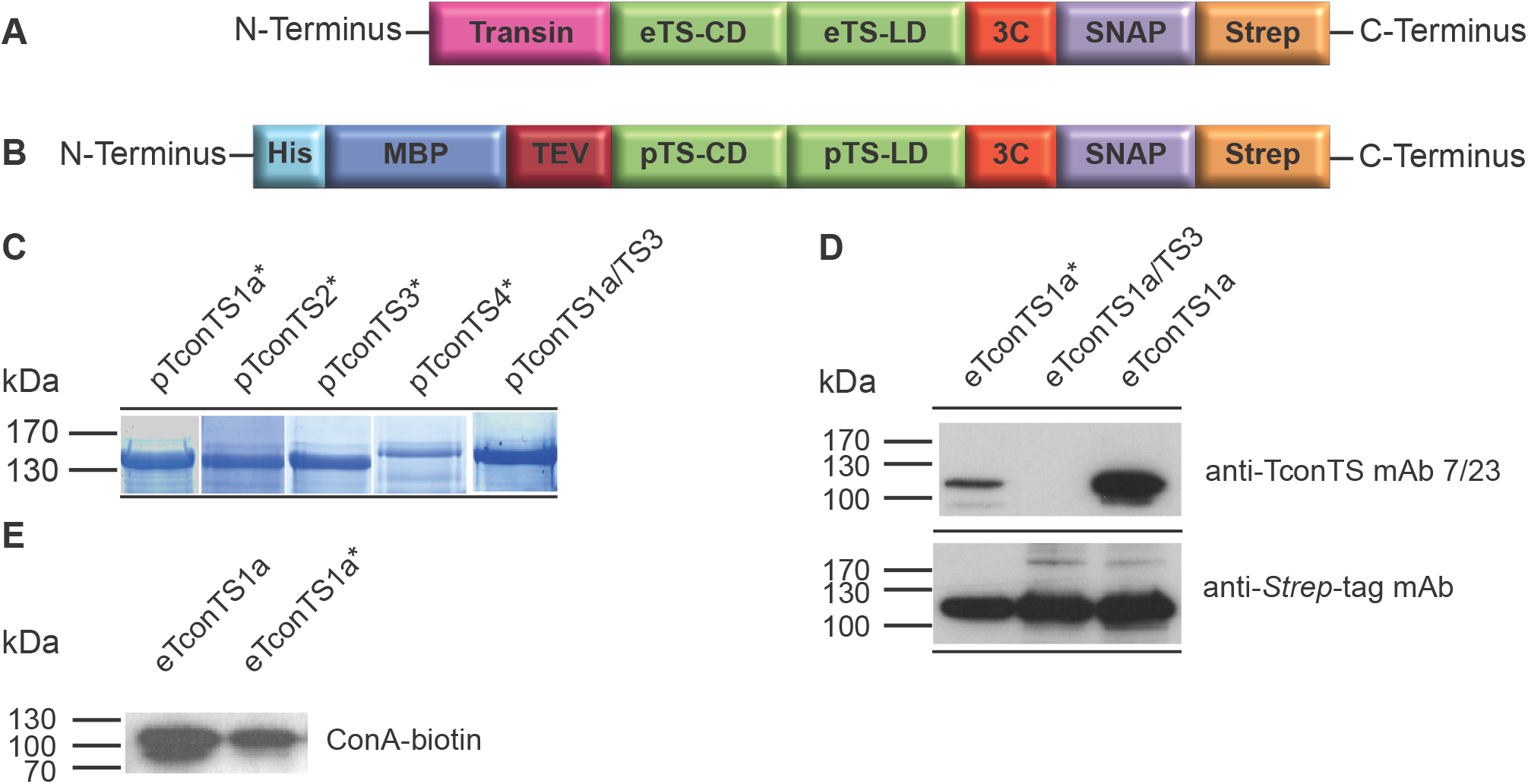
Recombinant TconTS constructs generated for expression in *E. coli* and CHO-Lec1 cells. **A-B:** Schematic presentation of recombinant TconTS constructs for expression in CHO-Lec1 fibroblasts eTconTS (**A**) and *E.coli* pTconTS (**B**). Fusion tags flanking TconTS are: Transin: translocation signal peptide (only in **A**), His: poly histidine tag (only in **B**), MBP: maltose binding protein tag (only in **B**), TEV: *tobacco etch virus* protease cleavage site (only in **B**), 3C: human rhinovirus 3C protease cleavage site, SNAP: SNAP-tag, *Strep:* Strep-tag. **C:** SDS-PAGE of purified pTconTS constructs. 1-2 μg of protein were loaded as indicated on a 10 % SDS polyacrylamide gel, which was stained with Coomassie Brilliant Blue after electrophoresis. pTconTS constructs with Eco105I are indicated by *. Lanes 5 and 6 comprise domain swapped proteins as labelled (CD/LD). **D:** Western blot analysis of eTconTS constructs expressed in CHO-Lec1 fibroblasts. 100 ng of each eTconTS construct was used and detection was done employing anti-*Strep*-tag mAb and anti-TconTS mAb 7/23 as indicated (described under Methods). **E:** Lectin blot of eTconTS1a and eTconTS1a* using biotin-conjugated concanavalin A (ConA) lectin. 100 ng of enzyme were used, respectively. Detection was done using horseradish peroxidase-conjugated avidin-biotin-detection system (ABC-kit) from VectaShield.

**Table 4:** Possible domain swapped TconTS constructs

| Construct number | Catalytic domain | Lectin domain | Swap construct |
| --- | --- | --- | --- |
| 1 | TconTS1a | TconTS1a | TconTS1a* |
| 2 | TconTS2 | TconTS2 | TconTS2* |
| 3 | TconTS3 | TconTS3 | TconTS3* |
| 4 | TconTS4 | TconTS4 | TconTS4* |
| 5 | TconTS1a | TconTS2 | TconTS1a/TS2 |
| 6 | TconTS1a | TconTS3 | TconTS1a/TS3 |
| 7 | TconTS1a | TconTS4 | TconTS1a/TS4 |
| 8 | TconTS2 | TconTS1a | TconTS2/TS1a |
| 9 | TconTS2 | TconTS3 | TconTS2/TS3 |
| 10 | TconTS2 | TconTS4 | TconTS2/TS4 |
| 11 | TconTS3 | TconTS1a | TconTS3/TS1a |
| 12 | TconTS3 | TconTS2 | TconTS3/TS2 |
| 13 | TconTS3 | TconTS4 | TconTS3/TS4 |
| 14 | TconTS4 | TconTS1a | TconTS4/TS1a |
| 15 | TconTS4 | TconTS2 | TconTS4/TS2 |
| 16 | TconTS4 | TconTS3 | TconTS4/TS3 |

\* Mutated TconTS containing the inserted *Eco*105I restriction site

For example, TconTS3-LD might be able to modulate the activity of TconTS1-CD in the domain swapped fusion protein TconTS1a/TS3. Our choice of domain swap TconTS1-CD and TconTS3-LD was based on the following thoughts: (I) Although TconTS1 has a (>60-fold) higher specific activity for Sia transfer compared to hydrolysis, its sialidase activity can be well detected; whereas (II) for TconTS3 no sialidase activity could be detected under conditions used (Gbem et al. 2013). (III) Thus, fusing the LD of TconTS3 to the CD of TconTS1a (Table 4, construct number 6 TconTS1a/TS3) might modulate (increase or decrease) the ratio of Sia transfer to sialidase activities. Along this line, domain swapped pTconTS1a/TS3 (Table 4, construct 6) was expressed and purified using the same conditions described for expressing pTconTS* as well as wild type pTconTS (see Methods). After tandem affinity purification a major band close to 130 kDa was observed on SDS-PAGE gel (Figure 4C).

Besides bacterial expression, domain swapped eTconTS1a/TS3 was also expressed by CHO-Lec1 fibroblasts. When using the anti-*Strep*-tag antibody, a clear band for eTconTS1a/TS3, eTconTS1* and eTconTS1a at 120 kDa respectively can be seen in the corresponding western blot (Figure 4D). Lectin blot analysis using concanavalin A (ConA), a lectin specifically binding to mannose-glycans (Kalb & Levitzki 1968), revealed the presence of mannose containing glycan structures on eTconTS1a and eTconTS1a* (Figure 4E).

### Binding epitope determination of the monoclonal anti-TconTS mAb 7/23

In our previous study we have shown that the monoclonal anti-TconTS antibody (anti-TconTS mAb 7/23) does specifically recognise recombinant TconTS1 but not TconTS3 (Gbem et al. 2013). This specificity of the anti-TconTS mAb 7/23 can be a helpful tool in order to characterise domain swapped recombinant TconTS1a/TS3 and other swap constructs. The resulting question was whether the binding epitope of the antibody is located within the CD or LD.

To answer this question, different truncated recombinant pTconTS1a fragments were designed (Figure 5A). In total, five pTconTS1-CD (Figure 5A, constructs 1-5) and two pTconTS1-LD (Figure 5A, constructs 6 and 7) fragments, varying in length and containing an C-terminal *Strep-tag*, were cloned and expressed by *E. coli* Rosetta pLacI as described under Methods. Bacterial lysates containing truncated recombinant pTconTS fragments were used for epitope mapping, employing Western blot analysis using anti-*Strep*-tag antibody recognising the C-terminus and anti-TconTS mAb 7/23 (Figure 5B-D).

**Figure 5:**
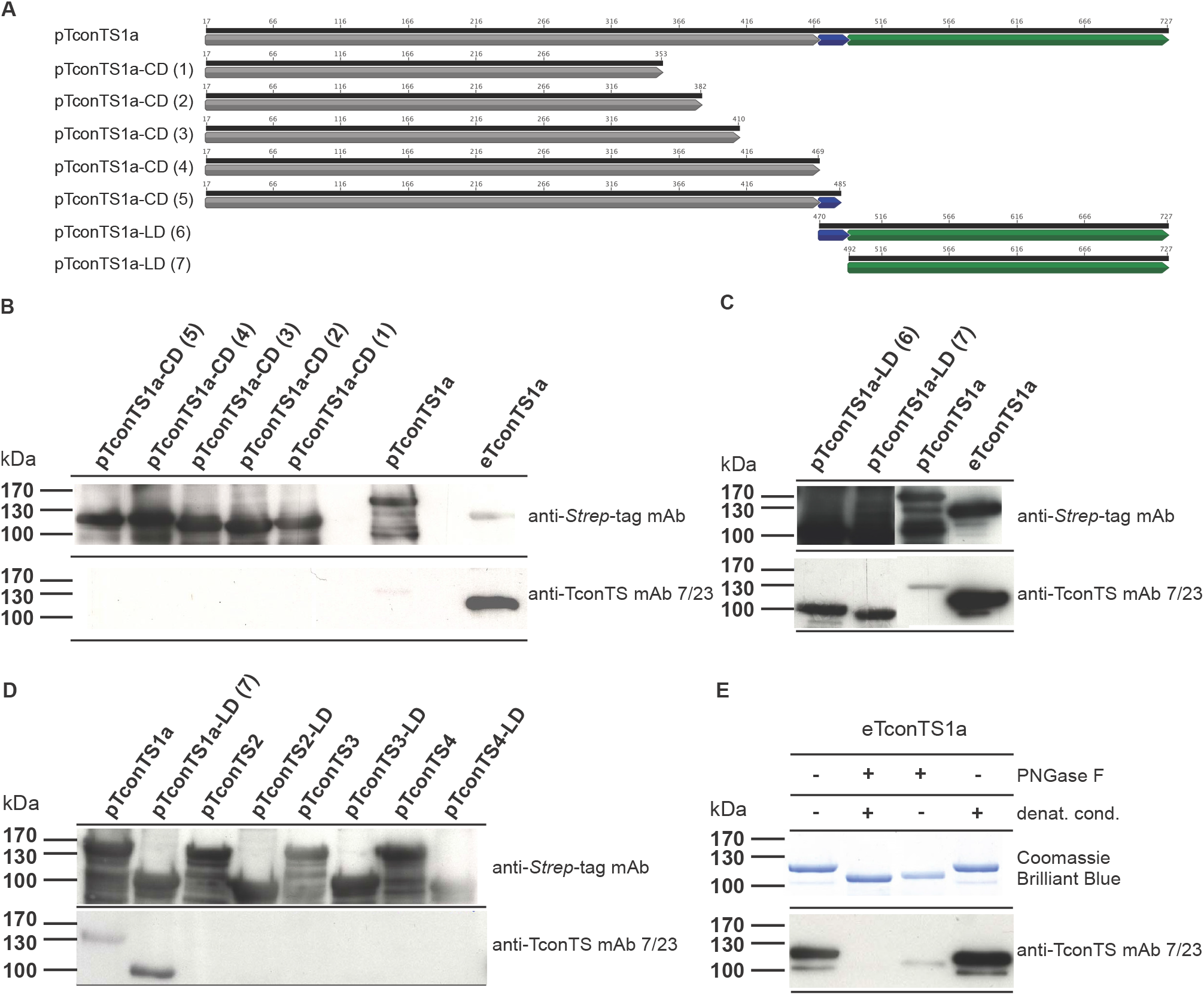
Epitope mapping of anti-TconTS mAb 7/23 binding epitope. **A:** Schematic presentation of wild type and truncated pTconTS1a constructs (1-5, CD: catalytic domain, 6-7 LD: lectin domain) used for anti-TconTS mAb 7/23 epitope mapping. Structural elements, such as catalytic domain (CD), α-helix and lectin domain (LD) are labelled in grey, blue and green respectively. **B-D:** Western blot analysis of pTconTS1a constructs (1-7) using anti-TconTS mAb 7/23 and anti-*Strep*-tag mAb as indicated (details under Methods). All pTconTS1a constructs were expressed in *E. coli* Rosetta pLacI and bacterial lysates were used for SDS polyacrylamide gel electrophorese. 100 ng eTconTS1a expressed and purified from CHO-Lec1 cells was used as a control in Western blot experiments. **E:** PNGaseF treatment of purified recombinant eTconTS1a. Recombinant eTconTS1a was deglycosylated using PNGaseF under native or denaturing conditions (denat. cond.) as described under Methods. 1 μg of eTconTS1a was used in SDS-PAGE analysis with subsequent Coomassie Brilliant Blue staining and 100 ng in Western blots using anti-TconTS mAb 7/23 and anti-*Strep*-tag for detection as indicated (details under Methods).

For all of the bacterial expressed constructs bands of similar intensities were detected by the anti-*Strep*-tag antibody at the expected molecular masses indicating that all recombinant proteins were synthesised by the bacteria at comparable levels. Notably, both pTconTS1a-LD constructs 6 and 7, were also detected by the anti-TconTS mAb 7/23 (Figure 5C) indicating that the binding epitope for mAb 7/23 is located in the LD. In contrast, none of the truncated recombinant pTconTS1-CD constructs (1-5) were recognised by the anti-TconTS mAb 7/23 (Figure 5B). Furthermore, when testing the other recombinant pTconTS2, pTconTS3 and pTconTS4 as well as their LDs (Figure 5D), none of them was recognised by the anti-TconTS mAb 7/23.

Interestingly, pTconT1a showed significantly lower signal intensity relative to eTconTS1a expressed by fibroblasts (Figure 5B). One of the major differences between proteins expressed in eukaryotic cells and those in prokaryotic cells is the lack of *N*-glycosylation in bacteria. To investigate the possible influence on anti-TconTS mAb 7/23 antibody binding to eTconTS1a, purified recombinant eTconTS1 was treated with peptide-*N*-glycosidase F (PNGase F), which specifically removes *N*-glycans.

Indeed, it was found that *N*-deglycosylation of eTconTS1a under native conditions drastically reduced the binding of anti-TconTS mAb 7/23 to the enzyme, indicated by the relatively weak bands at 110 kDa (Figure 5E). Furthermore, PNGaseF treatment under denaturing conditions completely eliminated the binding of anti-TconTS mAb 7/23 to eTconTS1a (Figure 5E). These observations indicate the involvement of *N*-glycan structures in the binding epitope of the anti-TconTS mAb 7/23 antibody binding to eTconTS1a.

### Activity of domain swapped TconTS1a/TS3

The crucial question of this study was which impact replacing the LD of TconTS1 with that from TconTS3 would have on the enzymatic activity of the fusion protein TconTS1a/TS3 (Table 4). To address this question, we compared the specific catalytic activities of wild type TconTS expressed by bacteria or fibroblasts using fetuin and lactose as Sia donor and acceptor substrates respectively. Reaction products 3-sialyllactose (3’SL) as indicator for TS activity, and *N*-Acetylneuraminic acid (Neu5Ac), as measure for sialidase activity, were quantified (Figure 6 and Table 5). A concentration series of pTconTS1a using up to 3 μg of enzyme, revealed a linear increase in 3’SL production up to 400 pmol due to Sia transfer activity (Figure 6A). In addition, only up to 3.5 pmol free Neu5Ac were detected within the same samples indicating slight sialidase activity (Figure 6B). Hence, 1 μg of pTconTS1a was used in subsequent time dependent measurements (Figure 6C and D). 3’SL production was linear up to 600 pmol, which was reached after 120 min of incubation.

**Figure 6:**
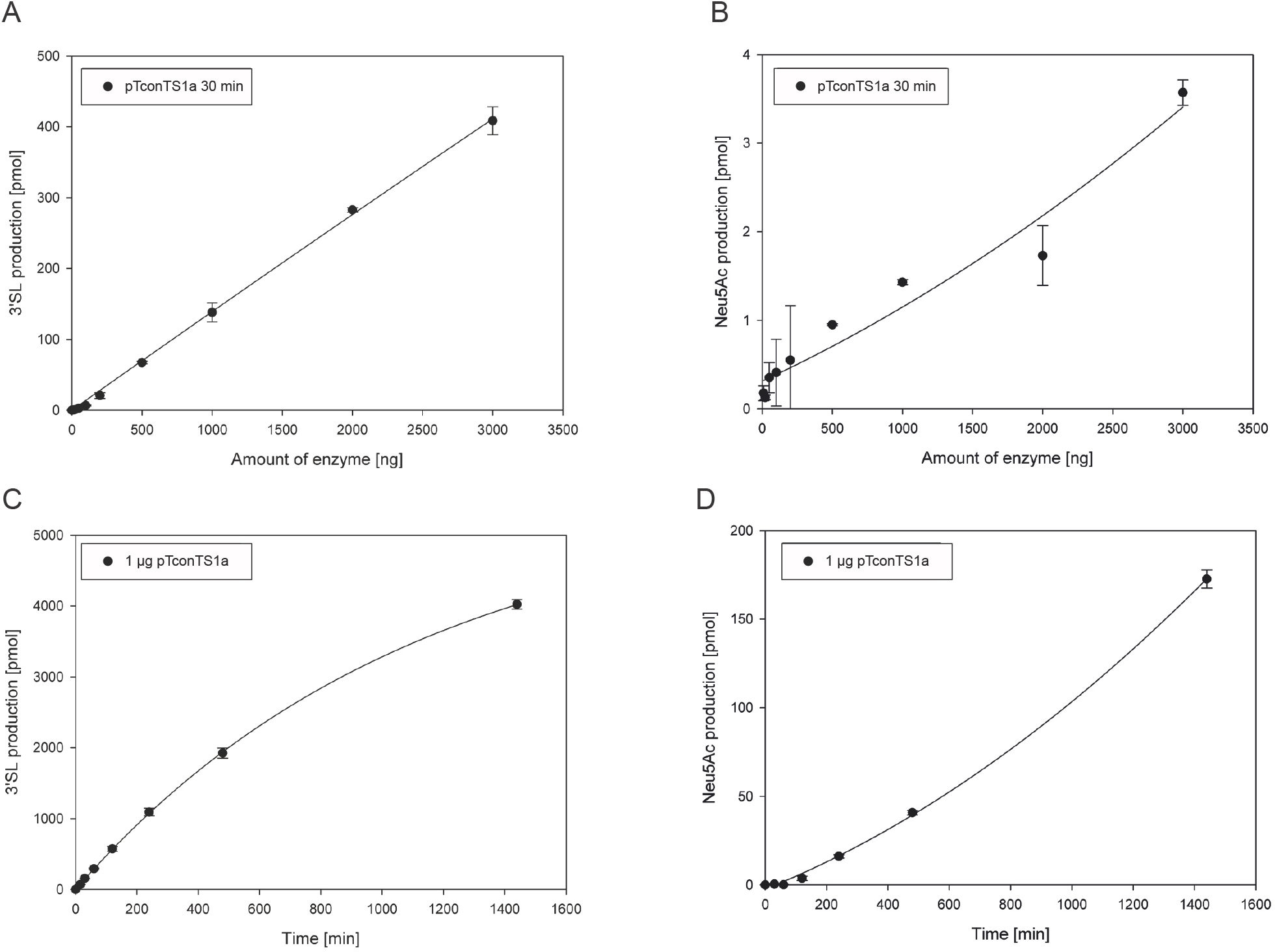
Enzymatic activities of bacterial expressed recombinant pTconTS1a. **A-B:** TconTS1a concentration dependent production of 3’SL (A) and Neu5Ac (B) using up to 3 μg of purified, bacterial expressed, recombinant TconTS (pTconTS). TS reactions were set up and analysed as described under Methods. Standard conditions with fetuin and lactose as Sia donor and acceptor substrates were incubated for 30 min at 37°C. **C-D:** Time dependency of TS and sialidase activities. Reactions were incubated for the indicated times ranging from 0-1440 min with 1 μg of purified pTconTS1a with standard fetuin and lactose concentrations (see Methods). Data points are means ± standard deviation of triplicates.

**Table 5:**
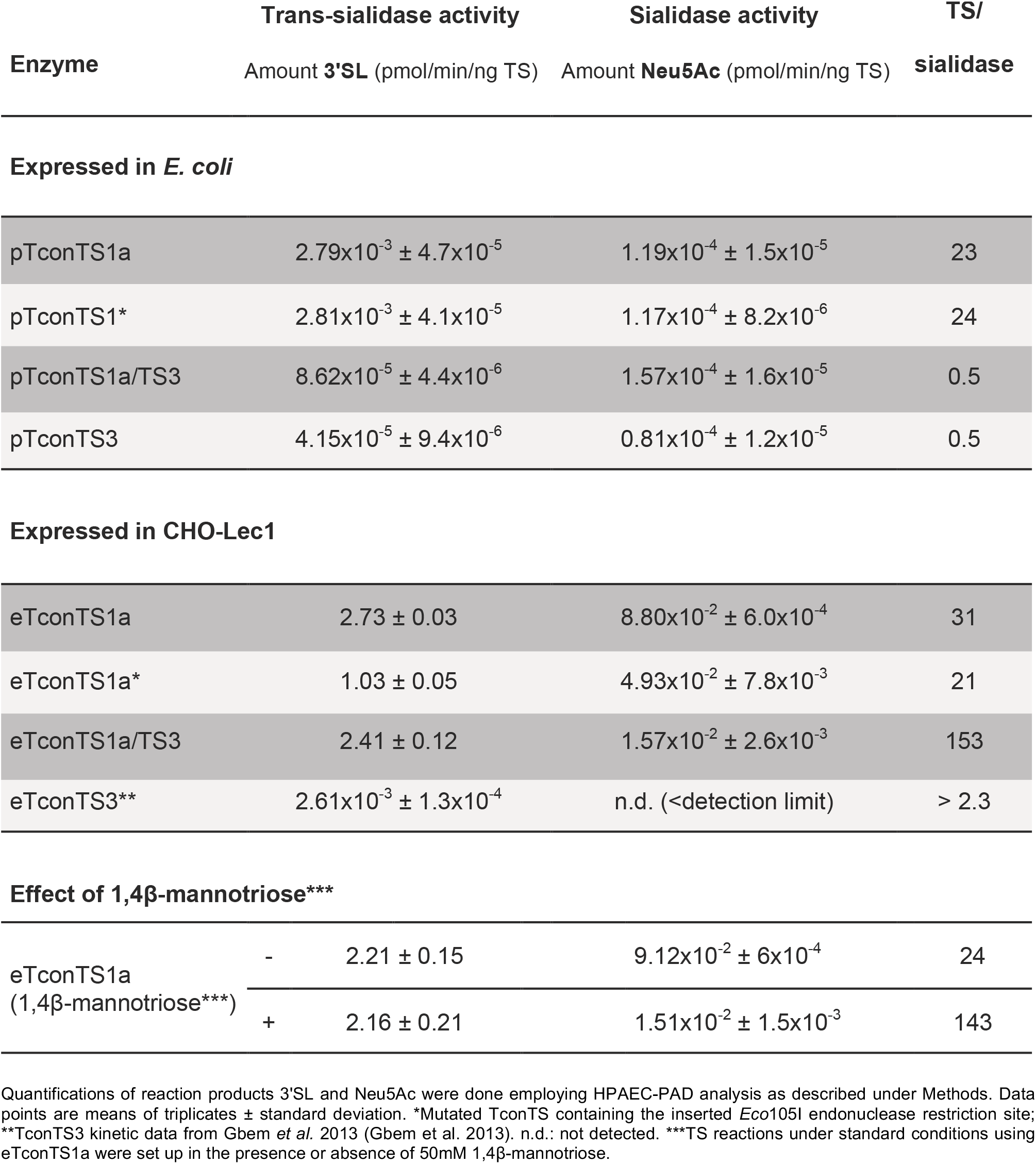
Specific catalytic activities of different TconTS enzymes expressed by *E. coli* and CHO-Lec1 cells also in the presence and absence of 5 mM 1-4β-mannotriose.

| Enzyme | Trans-sialidase activity | Sialidase activity | TS/ |
| --- | --- | --- | --- |
|  | Amount 3'SL (pmol/min/ng TS) | Amount Neu5Ac (pmol/min/ng TS) | sialidase |
| Expressed in *E. coli* |  |  |  |
| pTconTS1a | $2.79 \times 10^{-3} \pm 4.7 \times 10^{-5}$ | $1.19 \times 10^{-4} \pm 1.5 \times 10^{-5}$ | 23 |
| pTconTS1* | $2.81 \times 10^{-3} \pm 4.1 \times 10^{-5}$ | $1.17 \times 10^{-4} \pm 8.2 \times 10^{-6}$ | 24 |
| pTconTS1a/TS3 | $8.62 \times 10^{-5} \pm 4.4 \times 10^{-6}$ | $1.57 \times 10^{-4} \pm 1.6 \times 10^{-5}$ | 0.5 |
| pTconTS3 | $4.15 \times 10^{-5} \pm 9.4 \times 10^{-6}$ | $0.81 \times 10^{-4} \pm 1.2 \times 10^{-5}$ | 0.5 |
| Expressed in CHO-Lec1 |  |  |  |
| eTconTS1a | $2.73 \pm 0.03$ | $8.80 \times 10^{-2} \pm 6.0 \times 10^{-4}$ | 31 |
| eTconTS1a* | $1.03 \pm 0.05$ | $4.93 \times 10^{-2} \pm 7.8 \times 10^{-3}$ | 21 |
| eTconTS1a/TS3 | $2.41 \pm 0.12$ | $1.57 \times 10^{-2} \pm 2.6 \times 10^{-3}$ | 153 |
| eTconTS3** | $2.61 \times 10^{-3} \pm 1.3 \times 10^{-4}$ | n.d. (<detection limit) | > 2.3 |

Effect of 1,4β-mannotriose\*\*\*
|  |  |  |  |  |
| --- | --- | --- | --- | --- |
| eTconTS1a<br>(1,4β-mannotriose***) | - | $2.21 \pm 0.15$ | $9.12 \times 10^{-2} \pm 6 \times 10^{-4}$ | 24 |
| | + | $2.16 \pm 0.21$ | $1.51 \times 10^{-2} \pm 1.5 \times 10^{-3}$ | 143 |

Quantifications of reaction products 3'SL and Neu5Ac were done employing HPAEC-PAD analysis as described under Methods. Data points are means of triplicates ± standard deviation. \*Mutated TconTS containing the inserted *Eco*105I endonuclease restriction site; \*\*TconTS3 kinetic data from Gbem *et al.* 2013 (Gbem *et al.* 2013). n.d.: not detected. \*\*\*TS reactions under standard conditions using eTconTS1a were set up in the presence or absence of 50mM 1,4β-mannotriose.

For longer incubation times, a non-linear production of 3’SL was observed reaching a maximum of 4000 pmol 3’SL after 1440 min of incubation (Figure 6C). On the other hand, when looking at the release of Neu5Ac during the first 120 min hardly any free Neu5Ac was detected. Only after 120 min of incubation a non-linear increase of Neu5Ac production was observed resulting in 170 pmol free Neu5Ac after 1440 min (Figure 6D). The delayed increase in Neu5Ac production after 120 min of incubation indicates that pTconTS1a does not release free Neu5Ac from fetuin but rather from the reaction product 3’SL. Along this line, when considering the TS over sialidase activity (TS/sialidase) ratio pTconTS1a shows clear preference towards Sia-transfer to lactose (Table 5).

However, compared to catalytic activities observed for TconTS1 expressed by CHO-Lec1 fibroblasts (Koliwer-Brandl et al. 2011; Gbem et al. 2013), these findings clearly demonstrated that pTconTS1a exhibit around one order of magnitude lower overall enzymatic activity (Table 5). Nevertheless, for both TconTS1a enzymes, expressed by bacteria or fibroblasts, same preferences of trans-sialidase over sialidase activity in a similar order of magnitude have been observed (Table 5).

pTconTS1a produced 2.79 x 10^-3^ ± 4.7 x 10^-5^ pmol 3’SL and 1.19 x 10^-4^ ± 1.5 x 10^-5^ pmol Neu5Ac (Table 5). In contrast, pTconTS3 produced only 4.15 x 10^-5^ ± 9.4 x 10^-6^ pmol 3’SL and 0.81 x 10^-4^ ± 1.2 x 10^-5^ pmol Neu5Ac, indicating a low TS/sialidase ratio. Domain swapped pTconTS1a/TS3 exhibited different overall enzymatic behaviour compared to pTconTS1a, indicated by the formation of only 8.62 x 10^-5^ ± 4.4 x 10^-6^ pmol 3’SL and 1.57 x 10^-4^ ± 1.6 x 10^-5^ pmol Neu5Ac (Table 5). Interestingly, the TS/sialidase ratio of pTconTS1a/TS3 has shifted towards sialidase activity relative to that of pTconTS1a and is similar to that of pTconTS3 (Table 5). Modified pTconTS1* carrying the endonuclease restriction site Eco105I at the selected loop region following the domain-connecting α-helix (Figure 3) showed similar enzymatic activity as well as TS/sialidase ratio as pTconTS1a (Table 5).

Enzyme activity of CHO-Lec1 expressed eTconTS1a/TS3 was also investigated. Enzyme reactions were set up according to standard conditions using 50 ng of eTconTS1a/TS3 as well as eTconTS1a and eTconTS1a* as described under Methods. Results are summarised in Table 5. eTconTS1a produced about 2.73 ± 0.03 pmol 3’SL and 8.80 x 10^-2^ ± 6.0 x 10^-4^ pmol Neu5Ac. In contrast, domain swapped eTconTS1a/TS3 produced comparable amounts 2.41 ± 0.12 pmol of 3’SL but only 1.57 x 10^-2^ ± 2.6 x 10^-3^ pmol of Neu5Ac. In that respect, it can be seen that the TS/sialidase ratio has drastically shifted towards TS activity in case of eTconTS1a/TS3 relative to that of eTconTS1a (Table 5). This change in enzymatic behaviour is the exact opposite of what has been observed for pTconTS homologues. TS and sialidase activity of eTconTS1a* indicated by the production of 1.03 ± 0.05 pmol 3’SL and 4.93 x 10^-2^ ± 7.8 x 10^-3^ pmol Neu5Ac respectively, are both lower than that of eTconTS1a. However, TS/sialidase ratio is similar for both enzymes.

We have previously demonstrated that TconTS1a-LD is a carbohydrate binding domain with a specific affinity for oligomannose oligosaccharides (Waespy et al. 2015). Hence it was hypothesised that the LD of TconTS may bind to the same substrate glycan as its CD, but at a different position. This would provide a mechanism how interaction of the LD could directly influence the overall enzyme activity. To investigate this hypothesis, eTconTS1a was incubated with fetuin and lactose as Sia donor and acceptor substrates under standard conditions, in the presence and absence of 5 mM 1,4β-mannotriose and the reaction products 3’SL and Neu5Ac were analysed (described under Methods). The presence of 1,4β-mannotriose did not have a significant effect on the production of 3’SL (Table 5). Interestingly, in the presence of 1,4β-mannotriose the release of free Neu5Ac was about 6-fold lower than without this trisaccharide, resulting in a corresponding increase in the ratio of TS over sialidase activity (Table 5). It is important to point out here that this effect of 1,4β-mannotriose is in a similar range to that obtained for the domain swapped eTconTS1a/TS3 relative to eTconTS1a, where the increase of TS/sialidase ratio is about 4,9 (Table 5).

## Discussion

Based on our findings, namely identifying TconTS-LD as a carbohydrate-binding domain with specific affinities to oligomannosyl oligosaccharides (Waespy et al. 2015), we have hypothesised that the LD modulates TconTS catalytic activities, due to additional binding of TconTS-LD to the same substrate, influencing the overall binding affinities or catalytic turnover. To investigate the hypothesis of a cooperative binding of TS-CD and LD to the same substrate, we established a strategy for a modular recombination of TconTS-CD and LD allowing us to efficiently swap CD and LD between different TconTS. As a proof of principle, the CD of the highly active TconTS1a was fused to the LD of TconTS3, a comparatively less active TS (Gbem et al. 2013). Furthermore, TconTS1a-LD binds to 1,4β-mannotriose, whereas TconTS3-LD did not bind to this oligosaccharide (Waespy et al. 2015). Therefore, we investigated the impact of 1,4β-mannotriose on enzymatic activity of recombinant TconTS. Notably, in the presence of this trisaccharide the catalytic activity of eTconTS1a was modulated resembling the activity of the domain swap eTconTS1a/TS3. These results can be explained by cooperativity between CD and LD in TconTS1a, which catalytic activity is modulated by oligosaccharide binding to its LD as discussed below.

*In silico structural insights into the contact site between CD and LD of trypanosomal TS* Buschiazzo and co-worker first observed that the interface between TcruTS-CD and LD is significantly larger (2550 Å^2^) compared to other bacterial and viral sialidases (1300 – 1600 Å^2^) (Buschiazzo et al. 2002). Furthermore, Amaya and co-worker found that also the molecular surface of the contact site in TranSA is more extended (about 2600 Å^2^) (Amaya et al. 2003). Along this line, we calculated the area of the molecular interfaces from our *in silico* models of TconTS1a, TconTS2, TconTS3, TconTS4 and domain swap TconTS1a/TS3 and observed area sizes analogous to those obtained for TcruTS and TranSA (Amaya et al. 2003) (Table 1). In conclusion, in both publications (Buschiazzo et al. 2002; Amaya et al. 2003) the authors predicted a relatively rigid overall TS core structure precluding a direct involvement of the LD in enzymatic catalysis. This is in agreement with our hypothesis that the LD is indirectly involved in enzyme activity instead, potentially by modulating the affinities of TS for several donor/acceptor substrates (Waespy et al. 2015). In conclusion, the extended interfaces between CD and LD seems to be a typical feature of trypanosomal TS. Therefore, we investigated the structural architecture at the contact sites between TconTS CD and LD in detail. Interestingly, our TconTS homology models revealed that the majority of amino acids localised at the contact site between CD and LD are well conserved among TconTS family members (Figure 1D, Table 3), indicating a high evolutionary pressure to maintain these critical amino acids in all TconTS (Gbem et al. 2013). When calculating the hydrogen bond network formed between residues at the interface between CD and LD, 11 to 13 potential hydrogen bonds were observed for TconTS1a through TconTS4, which is similar to the number found in TranSA (Amaya et al. 2003). This is in contrast to other sialidases, where only about half of the number have been found. Mainly two separate areas, 1 and 2, comprise the majority of hydrogen bonds formed, evenly distributed over both (Figure 1A). Area 1 is located at the region where CD, LD and the α-helix are in close contact (Figure 1A and B), whereas Area 2 is located more closely to the active site of CD (Figure 1A and C). We assume that the contacts at Area 2 keep the catalytic grove of CD (Figure 1A purple square) close to the hypothesised binding site of the LD (Waespy et al. 2015), thus providing a rigid overall structure of both domains relative to each other. Strikingly, amino acid residues involved in the hydrogen bond network formation at both sites, 1 and 2, are well conserved (Table 3). Since all these amino acid residues are highly conserved among TS family despite their different catalytic activities (e.g. of TranSA), it appears unlikely that their conservation is essential for a specific catalytic activity. Nevertheless, the proposed cooperative binding of CD and LD requires a stabilised conformation of the two domains as provided by the conserved hydrogen bond networks.

### The potential of swapping catalytic and lectin-like domains between different TS

Based on the fact that the majority of amino acid residues localised at the contact sites between CD and LD are well conserved in the TconTS family, we rationalised that it might be possible to swap the domains of different TconTS to investigate the influence of LD on enzyme activities. To determine *in silico* structural stability of domain swap TconTS a structure homology model was calculated, using TconTS1a/TS3 amino acid sequence (Figure 2). In this model TconTS1a-CD and TconTS3-LD exhibited similar structure topology to the corresponding holoenzymes, TconTS1a and TconTS3 (Figure 2). Interestingly, when investigating the interface between CD and LD of TconTS1a/TS3, 13 hydrogen bonds were observed formed by amino acid residues, which were predicted to be essential for conformation stability of the wild type TconTS as discussed above. These observations underline the possibility that such an rearrangement of TconTS genes has occurred during evolution, which would provide an explanation for the different phylogenetic relationships of CD and LD in TS from African trypanosomes (Gbem et al. 2013).

At first sight, the α-helix itself might present a potential target for the restriction site introduction, but amino acid sequence alignments of TconTS revealed that the majority of amino acid residues of the α-helix are well conserved (Figure 1D, Figure 3A and Table 3). Thus, the insertion or exchange of amino acids in the sequence of the α-helix is expected to interfere with structure and/or interactions (see above). However, right after this α-helix there is a hairpin loop between a disulphide bridge found in all TS (e.g. Cys493 and Cys503 in TconTS1a). This loop (Figure 3A) is highly flexible and does not contain any conserved motifs or structurally relevant elements. For us it appeared a suitable target for introducing a restriction site to swap CDs and LDs from different TconTS. Indeed, the homology model of TconTS1a/TS3 showed a similar overall structure compared to TconTS1a and TconTS3 (Figure 2), including a similar size of the molecular interface (Table 1). In addition, the model also revealed an extended hydrogen bond network between CD and LD of domain swapped TconTS1a/TS3 (Table 2 and 3). In conclusion, this model suggests that such a recombinant protein with swapped domains would fold properly.

Therefore, we applied this strategy to replace the LD of TconTS1a with the LD of TconTS3 and express recombinant TconTS1a/TS3 in bacteria (*E. coli* Rosetta DE3 pLacI) and eukaryotic cells (CHO-Lec1). The production of soluble active enzyme confirmed that CD and LD from different TS can be fused without complete loss of proper folding. The insertion of the dipeptide YV (Figure 3) to facilitate domain swaps did not affect TS or sialidase activity of the recombinant protein expressed in bacterial or eukaryotic cells (Table 5). Most importantly, the TS/sialidase ratio of TconTS1a* is similar to that of TconTS1a in expression systems (Table 5).

The literature shows that TS were predominantly expressed in bacteria (Buschiazzo et al. 2002; Coustou et al. 2012) but also in eukaryotic cells (Coustou et al. 2012; Koliwer-Brandl et al. 2011; Gbem et al. 2013). Therefore, we decided to expressed domain swapped TconTS1a/TS3 in both systems and to compare their activities. In this respect, we observed that pTconTS exhibit three orders of magnitudes lower enzyme activities compared to that of eTconTS (Table 5), in agreement with previously published data (Koliwer-Brandl et al. 2011; Gbem et al. 2013). One explanation for these observations is improper folding of pTconTS. Along this line, it is important to note that the TS/sialidase ratio is almost identical between pTconTS1a and eTconTS1a. This is in agreement with the assumption that only a few pTconTS1a molecules are properly folded with catalytic activity identical to eTconTS1a and almost all pTconTS1a molecules are improperly folded and inactive. In contrast, for pTconTS3 a less drastic loss of TS activity was obtained. Furthermore, pTconTS3 shows a pronounced different enzymatic activity compared to eTconTS3, indicated by a shift of TS/sialidase ratio from >2.3 (eTconTS3) to 0.5 (pTconTS3). A similar but more drastic inversion of TS/sialidase ratio was observed for the domain swap TconTS1a/TS3 from 153 (eTconTS1a/TS3) to 0.5 (pTconTS1a/TS3).

In summary, these data demonstrate that folding of TconTS is major challenge in bacteria and probably does not assemble the protein in the native eukaryotic system. Thus, the expression of TconTS in bacteria is not a suitable strategy to investigate structure-function relationship of these enzymes.

One major difference between proteins expressed in bacteria and eukaryotic cells is glycosylation on asparagine residues (*N*-glycosylation), which can affect the folding process as well as the activity of the protein. Along this line, *N*-glycosylation may play a more pivotal role in regulating TS-activity. For example, it has been reported that enzymatic deglycosylation of TvivTS1 expressed by *Pichia pastoris* led to a slight reduction in sialidase activity relative to the untreated TvivTS1 (Haynes et al. 2015).

The presence of several putative *N*-glycosylation sites in TconTS indicates that these enzymes contain *N*-glycans in both, CD and LD. Interestingly, the TconTS mAb 7/23, binds more strongly to eTconTS1a compared to pTconTS1a (Figure 5). It is unlikely that this is due to improper folding in the bacteria, since the anti-TconTS mAb 7/23 binds to the SDS-denatured protein in this experiment. However, it is likely that for efficient binding of mAb 7/23 to TconTS1-LD is supported by at least one *N*-glycan. Since PNGase treatment of eTconTS1a strongly reduced binding of this antibody (Figure 5). In this context it is important to note that native glycosylated TconTS was used for immunisation to generate this antibody (Tiralongo et al. 2003). Along this line, it must be keep in mind that the eukaryotic expression system (CHO-Lec1 cells) used in this and our previous studies (Koliwer-Brandl et al. 2011; Gbem et al. 2013) leads to high-mannose type *N*-glycans (Puthalakath et al. 1996) similar to those found on trypanosomal glycoproteins (Pontes de Carvalho et al. 1993), leading to the assumption of a similar situation for TconTS. It will be interesting to investigate the role of *N*-glycosylation of TconTS.

### Cooperativity between CD and LD of TconTS1

According to the assumption that the LD of TconTS indirectly influences enzyme activities, it was expected that LD of the less active TconTS3 may decrease the enzymatic activities, if attached to the CD of the more active TconTS1a. Surprisingly, replacement of the LD in eTconTS1a with eTconTS3-LD does not lead to a reduction of TS activity (Table 5). This indicates that eTconTS3-LD has no negative effect on trans-sialylation efficiency of eTconTS1a-CD. However, sialidase activity of eTconTS1a/TS3 was reduced by 82% compared to wild type eTconTS1a (Table 5), leading to a five-fold higher TS/sialidase ratio of eTconTS1a/TS3 relative to that of wild type eTconTS1a. It can be concluded that TconTS3-LD suppressed sialidase activity in eTconTS1a/TconTS3 relative to TconTS1a-LD.

Previously it was demonstrated that TconTS1a-LD binds to oligomannose trisaccharides, whereas no such interaction was observed for TconTS3-LD (Waespy et al. 2015). This raised the question whether occupation of the carbohydrate binding site in TconTS1a-LD modulates the enzymatic reaction at the active site of CD of TconTS1a. Therefore, we investigated whether 1,4β-mannotriose, one of the potential binding partners of TconTS-LD (Waespy et al. 2015), can influence the activities of eTconTS1a. Our results clearly demonstrated that in the presence of 5 mM 1,4β-mannotriose TS/sialidase ratio was increased more than five-fold, due to a suppression in sialidase activity (Table 5). Strikingly, this resembles almost precisely the effect of replacing the LD of TconTS1a with TconTS3-LD. Considering the diversity of TS in African trypanosomes, it will be interesting to generate also other domain swapped TconTS constructs for a deeper understanding of the interplay between the different CD and LD of trypanosomal TS. Preliminary data point also to a pivotal role of *N*-glycosylation of TconTS on its enzyme activities, encouraging investigations on *N*-glycosylation of TconTS and its impact on TS activity and consequently the impact on pathogenesis of trypanosomiasis.

We propose that structural architecture and orientation of the carbohydrate binding-site in TconTS-LD provides the possibility of multivalent ligands such as neighbouring cell surface glycoconjugates to bind to the LD with corresponding effects on the supramolecular arrangement of the glycocalyx components. Furthermore, the distance between the proposed binding site in the LD and the active site of the CD could allow cooperative interactions with both CD and LD.

For example it has been reported that glutamic acid/alanine-rich protein (GARP), a *T. congolense* stage specific glycoprotein, was co-purified with TS-form 1 but not TS-form 2, both isolated from *T. congolense* procyclic cultures (Tiralongo et al. 2003). Interestingly, for TS-form 1 significantly higher TS/sialidase ratio was observed, whereas relative sialidase activity was higher in TS-form 2, although for both preparations, TS-form 1 and TS-form 2, the same donor and acceptor substrate preferences were described (Tiralongo et al. 2003). Furthermore, GARP is glycosylated with high-mannose type and galactosyl oligosaccharides (Thomson et al. 2002) and is shown to be sialylated in TS reactions (Engstler et al. 1995). Together, these findings provide strong evidence for its multivalent binding potential to TconTS-CD and LD, as discussed earlier (Waespy et al. 2015).

In summary, this study clearly demonstrates the cooperative binding of CD and LD and the influence of LD on TconTS enzymatic activities. However, direct competition of 1,4β-mannotriose for the Sia acceptor binding-site in TconTS-CD can be excluded, since TS activity is not altered (Table 5). The discovery of this influence of LD on TconTS enzymatic activities provides novel insight into the complexity of TS catalytic mechanisms by demonstrating the modulatory effect of the LD and its interaction with glycan structures abundant on the surface of trypanosomes.

Furthermore, our results obtained from domain swapped TconTS1a/TS3 (Table 5) demonstrating the effect of LD modulating TS/sialidase ratio of TconTS now provide an explanation for the observations made for native TS from *T. congolense* by Tiralongo *et al*. (Tiralongo et al. 2003). Unravelling the roles played by glycans interacting with TS-LD and its modulation of TS activity opens new perspectives, not only for a better understanding of their mechanisms, but it also provides ideas how glycosylation can modulate other systems as well.

## Acknowledgments

We thank Nazila Isakovic for excellent technical assistance. We are thankful to Dr. Judith Weber for helpful discussions. We dedicate this work to Dr. Thaddeus T. Gbem who suddenly passed away September 2018, in Zaria/ Nigeria and his mentor Prof. Dr. Jonathan A. Nok who passed away in 2017. Dr. Gbem delivered key contributions to this paper and strongly supported writing the manuscript. Financial support of this project by the Deutsche Forschungsgemeinschaft (DFG; project grants to SK, Ke428/8–1, Ke428/10–1) and from Africa Centre of Excellence for Neglected Tropical Diseases and Forensic Biotechnology (ACE-NTDFB).

## Conflict of interest declaration

Authors declare to have no conflict of interests.

## Supplemental data

Table S1: List of primers used for cloning and Eco105I restriction site insertion.

## Experimental Section

### Materials

Unless stated otherwise, all chemicals and reagents used in this study were cell culture and analytical grade. Recombinant PNGaseF endoglycosidase was from New England Biolabs, United Kingdom. *Pfu* and *Taq* DNA polymerase, *Eco105*I, *Hind*III, *Nco*I, *Not*I, *Sal*I and *Spe*I Fast Digest restriction enzymes, T4-DNA ligase, isopropyl-β-D-1-thiogalactopyranoside (IPTG), Dithiothreitol (DTT), Coomassie Brilliant Blue (Page Blue), protein molecular weight marker (PageRuler), GeneJET DNA Gel Extraction Kit, BCA Protein Assay Kit, enhanced chemiluminescence system (ECL-Kit), Luria Broth (LB) microbial growth medium, were from Thermo Scientific, Germany. Biozym LE Agarose was from Biozyme Scientific, Germany. StrepTactin^®^ Sepharose^®^, purification buffers and *anti-Strep-tag*^®^ rabbit polyclonal antibody were from IBA, Germany. β-D-galactosyl-(1-4)-α-D-glucose (4α-lactose), *N*-acetyl-neuraminic acid (Neu5Ac), 3’sialyl-lactose (3’SL), β-D-glucopyranuronic acid (glucuronic acid), lyophilised Fetuin from fetal calf serum, polyethylene glycol sorbitan monolaurate (TWEEN^®^ 20), Ex-cell^®^ CD CHO media, PEI (Polyethylenimin) transfection reagent were from Sigma-Aldrich, Germany. Hygromycin and Gentamycin were purchased from PAA, Austria. 1-4β-D-mannotriose was from Megazyme, Ireland. Ultrafiltration units Vivacell and Vivaspin6 were from Sartorius, Germany. X-ray film were purchased from GE Healthcare, Sweden. Protino^®^ Ni-NTA Agarose and NucleoBond^®^ Midi Plasmid DNA Purification Kit were from Macherey-Nagel, Germany. Polyvinylidenedifluoride (PVDF) membranes were from Millipore, Germany. 96-well transparent microtitre plate were from Sarstedt, Germany. 6 mL gravity flow columns were from Biorad, Germany.

### Cloning of TconTS into modified pET28aMBP bacterial expression vector

DNA sequences encoding for TconTS1 (including truncated variants, Figure 5 A-G), TconTS2, TconTS3 and TconTS4 as well as their LDs were amplified from modified pDEF vector (Koliwer-Brandl et al. 2011; Gbem et al. 2013), using the corresponding set of sense and reverse primers listed in Table S1. The resulting PCR products were subcloned into modified pET28aMBP vector (Waespy et al. 2015) via *HindIII* and *Bam*HI (TconTS1, TconTS2 and TconTS3) or *Sal*I and *Bam*HI (TconTS4) following instructions of the manufacturers.

### Introduction of Eco105I restriction site into TconTS

To insert the *Eco*105I endonuclease restriction site into the appropriate location at the hairpin loop following the domain-connecting α-helix of TconTS, sense and reverse primers were designed annealing at this target location and comprising the *Eco*105I site (Table S1). DNA sequences coding for TconTS-CD-α-helix with the *Eco*105I site attached at the 5’-end were amplified using *HindIII* (TconTS1, TconTS2 and TconTS3) or *Sal*I (TconTS4) sense primer in combination with the corresponding *Eco*105I reverse primer (Table S1). In addition, DNA sequences encoding the corresponding TconTS-LD sequence with the *Eco*105I site attached to the 3’-end were amplified using *Bam*HI (TconTS1 through TconTS4) reverse and the appropriate *Eco*105I sense primer (Table S1). Both PCR products were digested using *Eco*105I Fast Digest restriction enzyme (Thermo Scientific, Germany), purified using GeneJET DNA Gel Extraction Kit (Thermo Scientific, Germany) and blunt-end ligated using T4-DNA ligase, following instructions of the manufacturers, to generate full TconTS sequence with the *Eco*105I restriction site inserted. Appropriate TconTS DNA sequences were cloned into modified pET28aMBP expression vector using *Bam*HI (TconTS1, TconTS2 and TconTS3) or *Sal*I (TconTS4) and *HindIII* restriction enzymes according to manufacturers instructions. All sequences and insertions were confirmed by DNA sequencing at the Max Planck Institute for Marine Microbiology, Bremen, Germany.

### Recombination of CDs and LDs from different TconTS

To generate domain swapped TconTS constructs, corresponding pET28aMBP plasmids encoding TconTS enzymes were digested with *Eco*105I and *HindIII* Fast Digest restriction enzymes (Thermo Scientific, Germany) to isolate the LD, which was subsequently cloned into a *Eco*105I and *HindIII* digested pET28aMBP plasmid coding for the CD of a TconTS variant different from that of the isolated LD, following the manufacturers instructions.

For expression of secreted TconTS constructs in mammalian fibroblasts, DNA sequences coding for mutated TconTS, only comprising the *Eco*105I endonuclease restriction site, and domain swapped TconTS constructs were subcloned into the modified pDEF expression vector using *Spe*I and *HindIII* restriction sites as described previously (Koliwer-Brandl et al. 2011).

### Purification of recombinant TconTS expressed by E. coli Rosetta (DE3) pLacI or CHO-Lec1 fibroblasts

Recombinant TconTS constructs were expressed by CHO-Lec1 fibroblasts or *E. coli* Rosetta (DE3) pLacI and subsequently purified as described previously (Waespy et al. 2015; Koliwer-Brandl et al. 2011).

In brief, for expression of TconTS constructs by *E. coli* Rosetta (DE3) pLacI, colonies freshly transformed with pET28aMBP plasmid, encoding the TconTS construct, were used for an overnight culture in Luria Broth (LB) medium containing 50 μg/mL kanamycin, incubated at 37°C and 240 rpm shaking. For large scale, 1 L of LB medium containing 50 μg/mL kanamycin was inoculated with 2 mL of the overnight culture and grown at 37°C and 240 rpm until an optical density of 0.5 at 600 nm was reached. Induction was done using Isopropyl-β-D-1-thiogalactopyranoside (IPTG, 0.1 mM final concentration) for 30 min at 37°C followed by an induction for 14 h at 4°C and 240 rpm. *E. coli* Rosetta (DE3) pLacI cultures expressing TconTS-LDs were induced with IPTG (0.1 mM final concentration) for 2 h at 37°C and 240 rpm. Purification of recombinant TconTS was done as described previously employing double affinity chromatography using Ni-NTA^®^ and *Strep-tag*^®^ chromatography (Waespy et al. 2015). Purified proteins were characterised by SDS-PAGE and Western blot analysis and quantified using BCA assay according to instructions of the manufacturers.

For stable transfection in CHO-Lec1 cells, DNA sequences coding for domain swapped TconTS1a/TconTS3 as well as sequences for TconTS1a* were subcloned into the mammalian expression vector pDEF. A schematic overview of the construct is shown in Figure 4A. Stable transfection, single clone selection, cultivation of TconTS expressing cells as well as purification of recombinant proteins was done as described previously (Koliwer-Brandl et al. 2011).

Recombinant TconTS constructs expressed by CHO-Lec1 fibroblasts were purified employing *Strep-*tag^®^ chromatography and characterised by SDS-PAGE and Western blot analysis as described previously (Koliwer-Brandl et al. 2011) and subsequently quantified by BCA assay. Cells transfected with TconTS1a* and TconTS1a/TS3 respectively, produced up to 5 mg/L of secreted protein in the cell culture supernatant directly after clonal selection.

### Trans-sialidase reactions of recombinant TconTS constructs expressed by E. coli and CHO-Lec1 fibroblasts

Purified recombinant TconTS enzymes were assayed for Sia transfer and sialidase activities using fetuin and lactose as Sia donor and acceptor substrates as described before (Koliwer-Brandl et al. 2011). In general, TconTS reactions in 50 μL reaction volume containing 10 mM phosphate buffer pH 7.4, the appropriate amount of recombinant TconTS enzyme (50 ng of TconTS expressed by CHO-Lec1 and 1 μg of enzyme expressed by *E. coli*) as well as fetuin (100 μg corresponding to 600 μM fetuin bound Sia) and lactose (2 mM final concentration) as Sia donor and acceptor substrates were incubated at 37°C for the times indicated. To determine the influence of 1-4β-mannotriose on enzyme activities of recombinant TconTS1a and TconTS2 expressed by CHO-Lec1 fibroblasts, 0.25 μmol (5 mM final concentration) of the trisaccharide were additionally added to the reaction mix described above. Reaction products 3’SL and Neu5Ac were quantified from chromatograms obtained, employing **h**igh **p**erformance **a**nion **e**xchange **c**hromatography with **p**ulsed **a**mperometric **d**etection (HPAEC-PAD) utilising a Dionex DX600 system in combination with a CarboPac PA100^®^ column (Dionex/ Thermo Scientific, Germany).

### Deglycosylation of TconTS using PNGaseF endogylcosidase

Purified TconTS expressed from CHO-Lec1 fibroblasts were deglycosylated using PNGaseF endoglycosidase under native and denaturing conditions according to instructions of the manufacturers (New England Biolabs, United Kingdom). In brief, for degylcosylation under denaturing conditions, 10 μg of TconTS enzyme in 20 μL of denaturing buffer (0.5 % SDS, 40 mM DTT) were incubated at 95°C for 10 min. After incubation, NP40 and Na2PO4 were added to final concentrations of 1 % (v/v) and 40 mM, pH 7.5, respectively. Finally, 2 units of PNGaseF endoglycosidase were added and the reaction mix was incubated at 37°C for 4 h. Deglycosylation reactions under native conditions were set up according to that under denaturing conditions but without incubation in denaturing buffer at 95°C for 10 min.

### SDS-PAGE, Western and Lectin Blot analysis

Protein samples were separated employing SDS-PAGE as described previously (Laemmli 1970) using a MiniProtean III electrophorese Unit (Bio-Rad, Germany) and stained with Coomassie Brilliant Blue (Thermo Scientific, Germany).

Western blot analysis was performed using primary rabbit anti-*Strep*-tag^®^ and mouse anti-TconTS mAb 7/23 for detection of recombinant TconTS as previously described (Koliwer-Brandl et al. 2011; Waespy et al. 2015). After blotting, membranes were developed using enhanced chemiluminescence system (ECL-Kit, Thermo Scientific, Germany) and X-ray film (GE Healthcare, Sweden) according to instructions of the manufacturers. For lectin blot analysis, 1 μg/mL biotinylated concanavalin A (ConA) lectin specifically recognising high-mannose type oligosaccharides was used to detect oligomannose N-glycans bound to TconTS. Detection was done using the horseradish peroxidase-based avidin-biotin-complex(ABC)-system from VECTASTAIN (1:40000 diluted) according to manufacturer instructions. X-ray films were developed in the same way as that for Western blot analysis.

### Homology Modelling and in silico calculations

Homology models of TconTS-LD containing and lacking the α-helix were calculated employing the molecular modelling software YASARA 13.3.26 (King & Sternberg 1996; Jones 1999; Mückstein et al. 2002; Canutescu et al. 2003; Qiu & Elber 2006; Krieger et al. 2009) as previously described (Koliwer-Brandl et al. 2011). In brief, crystal structure of *Trypanosoma cruzi* trans-sialidase (Buschiazzo et al. 2002) was used as a template structure (PDB: 3b69) for calculating the models. YASARA *homology modelling* module were modified manually from the default settings of the program: Modelling speed: slow, PsiBLASTs: 6, EValue Max: 0.5, Templates total: 1, Templates SameSeq: 1, OligoState: 4, alignments: 15, LoopSamples: 50, TermExtension:10. All homology models were energy minimized using the Molecular Dynamics module of YASARA with default settings. The molecular surface was calculated using the ESPPME (Electrostatic Potential by Particle Mesh Ewald) method of YASARA *Structure* with the following parameters: Force field: AMBER96 (Case et al. 2005), Algorithm used to calculate molecular surface: numeric, Radius of water probe: 1.4 Å, Grid solution: 3, Maximum ESP: 300 kJ/mol. Structural alignment of TconTS was generated pairwise based on structure using the MUSTANG (Multiple Structural Alignment Algorithm) module of YASARA *Structure* (Konagurthu et al. 2006).

Amino acid sequences alignments of TTS were performed employing the *Geneious Alignment* module of the software Geneious 5.5.5, using Blosum62 Cost Matrix (Kearse et al. 2012) with gap openings and extension 10 and 0.1 respectively. Adaptations and modifications were made using the same software. Increasing darkness of sheds indicates increasing number of identical amino acid residues at each position (black: 100 %; dark grey: 80 to 100 %; light grey 60 to 80 %; white: less then 60 % similarity). Numbers on top of each sequence indicate the corresponding residue number for the appropriate TTS sequence. Amino acid sequences of TconTS1a (CCD30508.1), TconTS2 (CCC91739.1), TconTS3 (CCD12651.1), TconTS4 (CCD12514.1), TcruTS (AAA66352.1), TbruTS (AAG32055.1), TranSA (AAC95493.1), TvivTS (CCD20961.1) were obtained from UniProt database.

## Notes

### Competing Interest Statement

The authors have declared no competing interest.

## References

Amaya, M.F. et al., 2003. The high resolution structures of free and inhibitor-bound *Trypanosoma rangeli* sialidase and its comparison with *T.cruzi* trans-sialidase. Journal of Molecular Biology, 325(4), pp.773–784.

Bayne, R.A. et al., 1993. A major surface antigen of procyclic stage *Trypanosoma congolense*. Molecular and biochemical parasitology, 61(2), pp.295–310.

Beecroft, R.P., Roditi, I. & Pearson, T.W., 1993. Identification and characterization of an acidic major surface glycoprotein from procyclic stage *Trypanosoma congolense*. Molecular and biochemical parasitology, 61(2), pp.285–294.

Buschiazzo, A. et al., 2000. Structural basis of sialyltransferase activity in trypanosomal sialidases. The EMBO journal, 19(1), pp.16–24.

Buschiazzo, A. et al., 2002. The crystal structure and mode of action of trans-sialidase, a key enzyme in *Trypanosoma cruzi* pathogenesis. Molecular cell, 10(4), pp.757–768.

Bütikofer, P. et al., 2002. Glycosylphosphatidylinositol-anchored surface molecules of *Trypanosoma congolense* insect forms are developmentally regulated in the tsetse fly. Molecular and biochemical parasitology, 119(1), pp.7–16.

Campetella, O.E. et al., 1994. A recombinant *Trypanosoma cruzi* trans-sialidase lacking the amino acid repeats retains the enzymatic activity. Molecular and biochemical parasitology, 64(2), pp.337–340.

Canutescu, A.A., Shelenkov, A.A. & Dunbrack, R.L., 2003. A graph-theory algorithm for rapid protein side-chain prediction. Protein science: a publication of the Protein Society, 12(9), pp.2001–2014.

Case, D.A. et al., 2005. The Amber biomolecular simulation programs. Journal of Computational Chemistry, 26(16), pp.1668–1688.

Clayton, J., 2010. Chagas disease 101. Nature, 465(7301), pp.S4–5.

Coustou, V. et al., 2012. Sialidases play a key role in infection and anaemia in *Trypanosoma congolense* animal trypanosomiasis. Cellular Microbiology, 14(3), pp.431–445.

Cremona, M.L. et al., 1995. A single tyrosine differentiates active and inactive *Trypanosoma cruzi* trans-sialidases. Gene, 160(1), pp.123–128.

Crennell, S. et al., 1994. Crystal structure of *Vibrio cholerae* neuraminidase reveals dual lectin-like domains in addition to the catalytic domain. Structure, 2(6), pp.535–544.

Engstler, M. & Schauer, R., 1993. Sialidases from African trypanosomes. Parasitology today (Personal ed.), 9(6), pp.222–225.

Engstler, M., Reuter, G. & Schauer, R., 1993. The developmentally regulated trans-sialidase from *Trypanosoma brucei* sialylates the procyclic acidic repetitive protein. Molecular and biochemical parasitology, 61(1), pp.1–13.

Engstler, M., Schauer, R. & Brun, R., 1995. Distribution of developmentally regulated trans-sialidases in the Kinetoplastida and characterization of a shed trans-sialidase activity from procyclic *Trypanosoma congolense*. Acta Tropica, 59(2), pp.117–129.

Gaskell, A., Crennell, S. & Taylor, G., 1995. The three domains of a bacterial sialidase: a beta-propeller, an immunoglobulin module and a galactose-binding jelly-roll. Structure, 3(11), pp.1197–1205.

Gbem, T.T. et al., 2013. Biochemical diversity in the *Trypanosoma congolense* trans-sialidase family. PLoS neglected tropical diseases, 7(12), p.e2549.

Haselhorst, T., 2004. NMR spectroscopic and molecular modeling investigations of the trans-sialidase from *Trypanosoma cruzi*. Glycobiology, 14(10), pp.895–907.

Haynes, C.L.F. et al., 2015. Production, purification and crystallization of a trans-sialidase from *Trypanosoma vivax*. Acta Cryst (2015). F71, 577–585 [doi:10.1107/S2053230X15002496], pp.1–9.

Jones, D.T., 1999. Protein secondary structure prediction based on position-specific scoring matrices. Journal of Molecular Biology, 292(2), pp.195–202.

Kalb, A.J. & Levitzki, A., 1968. Metal-binding sites of concanavalin A and their role in the binding of alpha-methyl d-glucopyranoside. The Biochemical journal, 109(4), pp.669–672.

Kamuanga, M., 2003. Socio-economic and cultural factors in the research and control of trypanosomiasis. PAAT Technical and Scientific series, (4), pp.1–67.

Kearse, M. et al., 2012. Geneious Basic: an integrated and extendable desktop software platform for the organization and analysis of sequence data. Bioinformatics (Oxford, England), 28(12), pp.1647–1649.

King, R.D. & Sternberg, M.J., 1996. Identification and application of the concepts important for accurate and reliable protein secondary structure prediction. Protein science: a publication of the Protein Society, 5(11), pp.2298–2310.

Koliwer-Brandl, H. et al., 2011. Biochemical characterization of trans-sialidase TS1 variants from *Trypanosoma congolense*. BMC biochemistry, 12(1), p.39.

Konagurthu, A.S. et al., 2006. MUSTANG: a multiple structural alignment algorithm. Proteins: Structure, Function, and Bioinformatics, 64(3), pp.559–574.

Krieger, E. et al., 2009. Improving physical realism, stereochemistry, and side-chain accuracy in homology modeling: Four approaches that performed well in CASP8. Proteins: Structure, Function, and Bioinformatics, 77 Suppl 9, pp.114–122.

Laemmli, U.K., 1970. Cleavage of structural proteins during the assembly of the head of bacteriophage T4. Nature, 227(5259), pp.680–685.

Luo, Y. et al., 1999. The 1.8 A structures of leech intramolecular trans-sialidase complexes: evidence of its enzymatic mechanism. Journal of Molecular Biology, 285(1), pp.323–332.

Montagna, G. et al., 2002. The trans-sialidase from the african trypanosome *Trypanosoma brucei*. European journal of biochemistry / FEBS, 269(12), pp.2941–2950.

Mückstein, U., Hofacker, I.L. & Stadler, P.F., 2002. Stochastic pairwise alignments. Bioinformatics (Oxford, England), 18 Suppl 2, pp.S153–60.

Nok, A.J. & Balogun, E.O., 2003. A bloodstream *Trypanosoma congolense* sialidase could be involved in anemia during experimental trypanosomiasis. Journal of biochemistry, 133(6), pp.725–730.

Oliveira, I.A. et al., 2014. Evidence of ternary complex formation in *Trypanosoma cruzi* trans-Sialidase Catalysis. Journal of Biological Chemistry, 289(1), pp.423–436.

Paris, G. et al., 2001. Probing molecular function of trypanosomal sialidases: single point mutations can change substrate specificity and increase hydrolytic activity. Glycobiology, 11(4), pp.305–311.

Pereira, M.E. et al., 1991. The *Trypanosoma cruzi* neuraminidase contains sequences similar to bacterial neuraminidases, YWTD repeats of the low density lipoprotein receptor, and type III modules of fibronectin. The Journal of experimental medicine, 174(1), pp.179–191.

Pontes de Carvalho, L.C. et al., 1993. Characterization of a novel trans-sialidase of *Trypanosoma brucei* procyclic trypomastigotes and identification of procyclin as the main sialic acid acceptor. The Journal of experimental medicine, 177(2), pp.465–474.

Puthalakath, H., Burke, J. & Gleeson, P.A., 1996. Glycosylation defect in Lec1 Chinese hamster ovary mutant is due to a point mutation in *N*-acetylglucosaminyltransferase I gene. The Journal of biological chemistry, 271(44), pp.27818–27822.

Qiu, J. & Elber, R., 2006. SSALN: an alignment algorithm using structure-dependent substitution matrices and gap penalties learned from structurally aligned protein pairs. Proteins: Structure, Function, and Bioinformatics, 62(4), pp.881–891.

Savage, A. et al., 1984. Structural studies on the major oligosaccharides in a variant surface glycoprotein of *Trypanosoma congolense*. Molecular and biochemical parasitology, 11, pp.309–328.

Schenkman, S. et al., 1991. A novel cell surface trans-sialidase of *Trypanosoma cruzi* generates a stage-specific epitope required for invasion of mammalian cells. Cell, 65(7), pp.1117–1125.

Schenkman, S. et al., 1994. A proteolytic fragment of *Trypanosoma cruzi* trans-sialidase lacking the carboxyl-terminal domain is active, monomeric, and generates antibodies that inhibit enzymatic activity. The Journal of biological chemistry, 269(11), pp.7970–7975.

Smith, L.E. & Eichinger, D., 1997. Directed mutagenesis of the *Trypanosoma cruzi* trans-sialidase enzyme identifies two domains involved in its sialyltransferase activity. Glycobiology, 7(3), pp.445–451.

Smith, L.E., Uemura, H. & Eichinger, D., 1996. Isolation and expression of an open reading frame encoding sialidase from *Trypanosoma rangeli*. Molecular and biochemical parasitology, 79(1), pp.21–33.

Thomson, L.M. et al., 2002. Partial structure of glutamic acid and alanine-rich protein, a major surface glycoprotein of the insect stages of *Trypanosoma congolense*. The Journal of biological chemistry, 277(50), pp.48899–48904.

Tiralongo, E. et al., 2003. Two trans-sialidase forms with different sialic acid transfer and sialidase activities from *Trypanosoma congolense*. The Journal of biological chemistry, 278(26), pp.23301–23310.

Waespy, M. et al., 2015. Carbohydrate recognition specificity of trans-sialidase lectin domain from *Trypanosoma congolense.* A. Acosta-Serrano, ed. PLoS neglected tropical diseases, 9(10), p.e0004120.

World Health Organization, 2013. Control and surveillance of human African trypanosomiasis. World Health Organization technical report series, (984), pp.1–237.

Zaccai, N.R. et al., 2003. Structure-guided design of sialic acid-based Siglec inhibitors and crystallographic analysis in complex with sialoadhesin. Structure, 11(5), pp.557–567.

Zamze, S.E. et al., 1990. Characterisation of the asparagine-linked oligosaccharides from *Trypanosoma brucei* type-I variant surface glycoproteins. European journal of biochemistry / FEBS, 187(3), pp.657–663.

Zamze, S.E. et al., 1991. Structural characterization of the asparagine-linked oligosaccharides from *Trypanosoma brucei* type II and type III variant surface glycoproteins. The Journal of biological chemistry, 266(30), pp.20244–20261.

