## Supplemental Table S1 for "Cooperativity of catalytic and lectin-like domain of *T. congolense* trans-sialidase modulates its catalytic activity"

### Supplemental data

**Table S1.** List of primers used for cloning and *Eco*105I restriction site insertion.

| Gene | Forward primer (restriction enzyme) | Reverse primer (restriction enzyme) |
|---|---|---|
| For bacterial expression by *E. coli* Rosetta (DE3) pLacI | | |
| TconTS1a | GCAAGCTTCAGTGCTGCGACCACATG (*Hind*III) | CGGGATCCGTCGCTCCCAGGCACACG (*Bam*HI) |
| TconTS2 | GCAAGCTTGCCCAGTGCATCTCAACG (*Hind*III) | GCGGATCCAGACACGGGATGCACATC (*Bam*HI) |
| TconTS2-LD | GCAAGCTTCTGGAGGATGAGATGGAGG (*Hind*III) | |
| TconTS3 | GCAAGCTTTCTGGAAACGGACGAACG (*Hind*III) | GCGGATCCGAGGTAAAGTGACTCCAGTTC (*Bam*HI) |
| TconTS3-LD | GCAAGCTTCTAGAAGACGAGCTGGAAAGC (*Hind*III) | |
| TconTS4 | GCGTCGACATCCTACAAGAAAGCTC (*Sal*I) | GCGGATCCCCTTGCTGCTCTTTTAAGTAAT (*Bam*HI) |
| TconTS4-LD | GCGTCGACCTCGCTGACGAACTGAAG (*Sal*I) | |
| Epitope mapping for anti TS mAb 7/23 | | |
| TconTS1a-CD (1) | GCAAGCTTCAGTGCTGCGACCACATG (*Hind*III) | CGGGATCCGTCACTCGATTCGAATATC (*Bam*HI) |
| TconTS1a-CD (2) | | CGGGATCCATCCGAGCTGCCAGGACCA (*Bam*HI) |
| TconTS1a-CD (3) | | CGGGATCCATCACGATAATACGAGCCCT (*Bam*HI) |

| | | |
|---|---|---|
| TconTS1a-CD (4) | | CGGGATCCGTCCTGTGCCTTCCA CACC (*Bam*HI) |
| TconTS1a-CD (5) | | CGGGATCCGTCCACAAGGCGCA CAAGG (*Bam*HI) |
| TconTS1a-LD (6) | GCAAGCTTGACGAGCTGAAAAGC (*Hind*III) | CGGGATCCGTCGCTCCCAGGCA CACG (*Bam*HI) |
| TconTS1a-LD (7) | GCAAGCTTAACTGCCTCCCGGGC (*Hind*III) | |
| *Eco*105I insertion for TconTS domain swap | | |
| TconTS1a*-CD (*Eco*105I) | GCAAGCTTCAGTGCTGCGACCAC ATG (*Hind*III) | GCTACGTAATCGCCCGGGAGGC (*Eco*105I) |
| TconTS1a*-LD (*Eco*105I) | GCTACGTAAAATATGATCCCGGG (*Eco*105I) | CGGGATCCGTCGCTCCCAGGCA CACG (*Bam*HI) |
| TconTS2*-CD (*Eco*105I) | GCAAGCTTGCCCAGTGCATCTCA ACG (*Hind*III) | GCTACGTATTTGTTCAGTTGAC (*Eco*105I) |
| TconTS2*-LD (*Eco*105I) | GCTACGTAAAGCGCCGGAGCGG C (*Eco*105I) | GCGGATCCAGACACGGGATGCA CATC (*Bam*HI) |
| TconTS3*-CD (*Eco*105I) | GCAAGCTTTCTGGAAACGGACGA ACG (*Hind*III) | GCTACGTAACCATCCGGTGAGG (*Eco*105I) |
| TconTS3*-LD (*Eco*105I) | GCTACGTAGATTATACTGAGGG (*Eco*105I) | GCGGATCCGAGGTAAAGTGACTC CAGTTC (*Bam*HI) |
| TconTS4*-CD (*Eco*105I) | GCGTCGACATCCTACAAGAAAGC TC (*Sal*I) | GCTACGTACCCGGTAGTTGCAG (*Eco*105I) |
| TconTS4*-LD (*Eco*105I) | GCTACGTAGATGGCAGCGATTGC (*Eco*105I) | GCGGATCCCCTTGCTGCTCTTTT AAGTAAT (*Bam*HI) |

*Mutated TconTS containing the inserted *Eco*105I restriction site. Endonuclease restriction sites are underlined.
